# QuantCell: machine learning-based cell annotation leveraging quantitative imaging profiles

**DOI:** 10.64898/2026.02.15.706033

**Authors:** Wade Boohar, Bowen Wang, Zachary Thomas, Anna Nogalska, Rong Lu

## Abstract

Spatial omics enables high-resolution, multiplexed in situ imaging of RNA and protein expression, but cell annotation remains challenging in complex tissues, particularly for rare, functionally important cell populations. We developed QuantCell, a machine learning framework that leverages quantitative imaging data to improve cell annotations derived from qualitative profiles. QuantCell evaluates multiple established supervised machine learning models to identify the best-performing model and applies a user-defined false discovery rate (FDR) to ensure high-confidence annotation. Using PhenoCycler imaging of mouse bone marrow, QuantCell increased the proportion of annotated cells from 41.0% to 94.4% at a 5% FDR, achieving 97.3% accuracy on a held-out test set. QuantCell achieved higher precision than the comparison methods on our bone marrow datasets and externally annotated colon and lymph node datasets from published studies. While comparison methods favored abundant cell populations, QuantCell maintained consistently high precision across all evaluated cell types. QuantCell provides an effective and flexible workflow compatible with diverse imaging platforms and adaptable to project-specific needs, enabling robust identification of both abundant and rare cell types in spatial omics studies.

## Introduction

Recent advances in spatial omics technologies have revolutionized our ability to investigate cellular organization and function within native tissue environments (Cheng *et al*, 2025; Lundberg & Borner, 2019; Marx, 2021; Seydel, 2025). These technologies generate high-resolution, spatially resolved molecular profiles that quantify gene or protein expression at a single-cell or subcellular level (Chen *et al*, 2022; Eng *et al*, 2019; Goltsev *et al*, 2018; Merritt *et al*, 2020; Stadler *et al*, 2013). A critical step in analyzing such data is to assign cell identities based on spatial molecular profiles, a process known as cell annotation or classification. This step sets a foundation for downstream analyses that infer cellular interactions, spatial organization, and functional states.

Several approaches have been developed to annotate cells based on spatial molecular profiles. Semi-supervised methods are widely used, relying on prior knowledge of marker genes (Coleman *et al*, 2023; Geuenich *et al*, 2021; Zhou *et al*, 2024) or predefined combinations of positive and negative markers for specific cell types (Ianevski *et al*, 2022; Mongia *et al*, 2024) to guide the annotation of cell clusters generated by unsupervised algorithms. In contrast, unsupervised methods cluster cells without prior knowledge or labels, and infer cell identities directly from the data (Kiselev *et al*, 2019; Kumagai, 2024; Marghi *et al*, 2024). More recently, supervised approaches have emerged for large-scale datasets, leveraging manual annotations of a subset of cells to train classifiers that subsequently annotate the remaining cells (Amitay *et al*, 2023; Shaban *et al*, 2024).

However, existing cell annotation methods face significant challenges when applied to complex tissues containing diverse cell populations with complex molecular markers. Rare cell types, such as stem cells and cancer stem cells, play pivotal roles in physiological and pathological contexts (Abbaszadegan *et al*, 2017; Altshuler *et al*, 2023; Nguyen *et al*, 2018; Patel *et al*, 2024), but are difficult to manually identify for training existing annotation models. The challenges are further exacerbated in spatial proteomics, where the number of measurable protein markers is constrained by cost and technical limits (Bressan *et al*, 2023; Wu *et al*, 2024). Yet, understanding protein expression and localization is essential for uncovering fundamental biological mechanisms and disease pathology (Blise *et al*, 2022; Hung & Link, 2011). Recent advances in artificial intelligence have shown great promise in addressing these challenges by leveraging the large-scale data generated by spatial imaging technologies (Bae *et al*, 2022; Dong & Zhang, 2022; Li *et al*, 2022). However, the available models and platforms are often not generalizable beyond their original experimental context, as their performance is highly optimized for a specific set of tissues, markers, and technical procedures (Ge *et al*, 2025).

In this study, we present QuantCell, a machine learning (ML) based framework designed to improve cell annotation using qualitative and quantitative features derived from multiplexed imaging. Here, “qualitative” refers to binary marker positivity calls (positive or negative for each protein marker in each cell), whereas “quantitative” refers to continuous intensity measurements across cellular compartments. In binary marker rule-based cell annotation, cell type identities are defined using combinations of binary marker patterns, similar to traditional immunophenotyping and flow cytometry approaches (Cossarizza *et al*, 2019). For example, in this paper, we define CD4 T cells as cells that are CD4^+^, CD3^+^, TCR^+^ and CD8^-^.

QuantCell integrates established ML methods into a practical, configurable workflow for extending marker-based cell annotations. First, it establishes a strategy for computationally generating training labels based on biologically defined marker combinations, eliminating the need for manual cell type annotation required by supervised methods, while enabling identification of rare but biologically important cell populations that are difficult to annotate manually. Second, QuantCell incorporates multiple summary statistics of marker intensities across each cellular compartment to improve predictive accuracy and robustness. Third, we implement cell type specific thresholding based on a user-defined false discovery rate (FDR) target to enhance annotation of rare cell populations that are often overlooked by conventional methods.

To demonstrate the utility of QuantCell, we imaged mouse bone marrow (BM) using the PhenoCycler (formerly CODEX) platform (Akoya Biosciences), which leverages DNA-barcoded antibodies for high-resolution, multiplexed imaging (Black *et al*, 2021). Images were processed using QuPath for cell segmentation, binary marker calling, and feature extraction (Bankhead *et al*, 2017; Loughrey *et al*, 2018), generating intensity-based summary statistics across imaging channels and cellular compartments. Mouse BM contains over 20 distinct cell types with a wide range of frequencies (Baccin *et al*, 2020; Bryder *et al*, 2006; Ding & Morrison, 2013; Dolgalev & Tikhonova, 2021; Nombela-Arrieta & Manz, 2017). Notably, hematopoietic stem and progenitor cells (HSPCs) in the BM are defined by complex combinations of cell surface markers (Purton, 2022; Spangrude *et al*, 1988) (Table EV1), making BM an ideal system for evaluating the accuracy and robustness of QuantCell. In addition to mouse BM, we further evaluated QuantCell using ground-truth data from published studies of colon and lymph node tissues (Schürch *et al*, 2020; Shaban *et al*, 2024) and compared its performance with existing cell annotation methods.

## Results

### QuantCell concept and design

We demonstrated the performance of QuantCell using data acquired from the PhenoCycler (Black *et al*, 2021) platform. Mouse BM sections were imaged after staining with barcoded antibodies targeting 25 distinct proteins that are established cell type markers (Fig. 1A, Appendix Figs S1 and S2). These markers were able to annotate 21 BM cell types (Table EV1), which collectively comprise the vast majority of cells in the BM. Following the PhenoCycler manufacturer’s recommendation, cell segmentation was performed using nuclear staining in QuPath (Bankhead *et al*, 2017), an open-source bioimage analysis software platform (Fig. 1B, Fig. EV1A). QuantCell performs with comparable precision using nuclei-based and membrane-based cell segmentation (Fig. EV1). Subsequently, a binary label (positive or negative) for each protein marker was assigned to each segmented cell using QuPath’s cell classifier, which requires manual qualitative validation of marker expression in only a few dozen cells (Fig. 1C).

**Figure 1.**
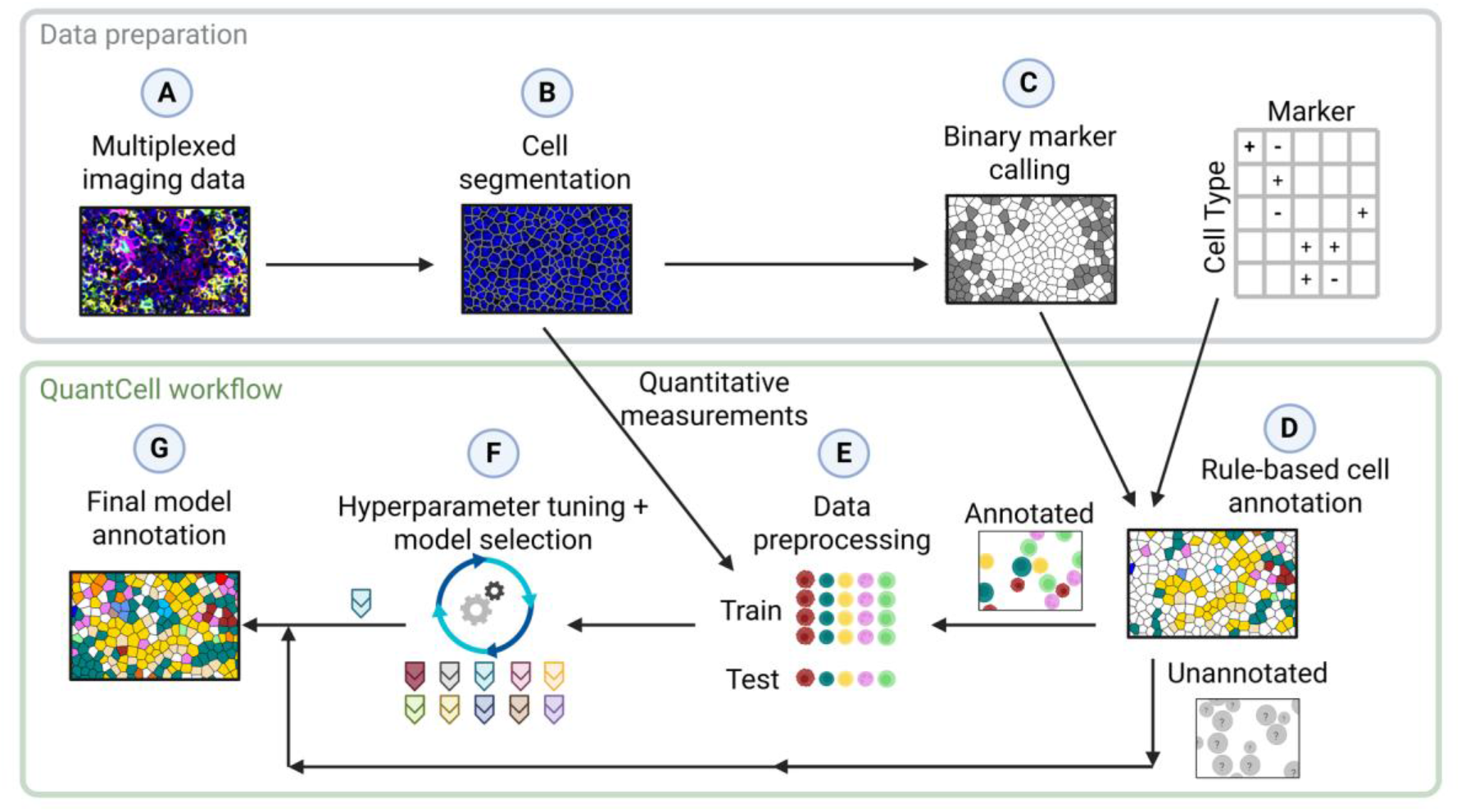
Overview of QuantCell annotation workflow. (A) Tissue sections are prepared, stained, and imaged. (B) Cell segmentation is performed using QuPath (Bankhead *et al*, 2017), based on 4′,6-diamidino-2-phenylindole (DAPI) nuclei staining. (C) Each protein marker is manually annotated as positive or negative in a small subset of cells. These annotations are then propagated to the remaining cells using QuPath. (D) Cells are classified based on manually defined, expected combinations of positive and negative markers for each cell type. (E) Quantitative measurements are generated by QuPath for each marker and cellular compartment, by summarizing the intensity distribution using the minimum, maximum, mean, median, and standard deviation. The resulting data are scaled and encoded. 80% of the cells annotated by the binary marker rule-based method are used for model selection and training, while the remaining 20% are reserved as a test dataset. (F) Multiple ML models are evaluated using 10-fold cross-validation after hyperparameter tuning. (G) The final model is applied to classify cells that are unannotated **in** (D), using a 5% FDR threshold.

Cell annotation in QuPath requires manually identifying representative cells for each cell type, which is labor-intensive, especially for rare cell types or those defined by multiple markers. QuantCell leverages binary marker profiles from QuPath to classify cells based on expected combinations of positive and negative markers for individual cell types (Fig. 1D, Table EV1). QuantCell does not rely on spatial context for cell annotation, enabling its application to tissues lacking fixed architecture, such as bone marrow, as well as to disease settings where spatial organization may be disrupted or variable. Using the binary marker rule-based approach based on qualitative marker expression profiles, 41.0% of BM cells were annotated.

To classify the remaining cells, QuantCell leverages quantitative imaging data to improve cell annotation derived from qualitative data. Using rule-based annotations as the ground truth, QuantCell trains ML models for cell classification on the quantitative marker expression measurements of individual cells (Fig. 1E). This process involves testing appropriate ML models and fine-tuning their hyperparameters to optimize performance (Fig. 1F). For model selection and training, 80% of the cells annotated by the rule-based approach were used, with the remaining 20% reserved as a test dataset. Model training, hyperparameter tuning, and model selection were performed using cross-validation within the training dataset only. The test dataset was used for final performance evaluation and to determine the FDR-based classification thresholds that were then applied to classify cells that were not annotated by the rule-based approach (Fig. 1G). For the accuracy evaluation in Fig. 3A, C and E, thresholds were selected on a separate validation set (see Methods).

### Selection and optimization of ML models for cell classification

QuantCell systematically compares a variety of classification models based on their macro average precision (AP) score (Figs 1F, 2A). The AP score is derived from the precision–recall curve (Fig. 2B), where precision reflects the accuracy of cell type prediction by the model, and recall represents the fraction of ground-truth cells that are correctly annotated. The AP score, which corresponds to the area under the precision–recall curve, provides a summary measure of model performance. Precision, recall, and AP are calculated separately for each cell type, and the unweighted average across all cell types defines the macro AP score.

**Figure 2.**
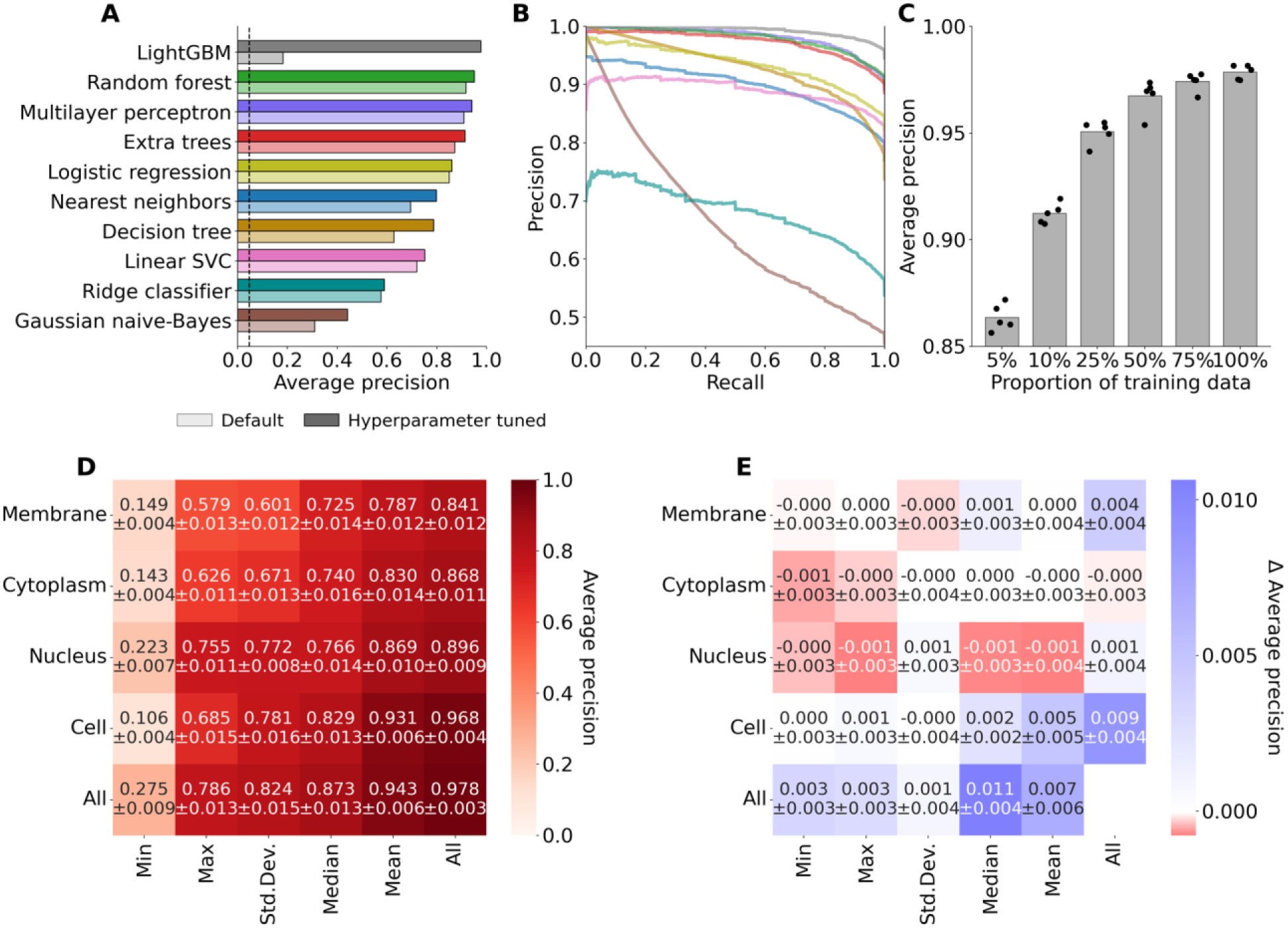
Selection and optimization of ML models for cell classification. (A) Macro AP score for each classifier with default and tuned hyperparameter settings. The dashed line indicates the random prediction baseline. Bars show the mean macro AP over 10-fold cross-validation (n = 10 folds); SDs are given in Table EV2. (B) Precision-recall curve for tuned classifiers. Data represent the unweighted average of all cell types. Curves are the mean over the 10 cross-validation folds. (C) Comparison of QuantCell performance using varying proportions of the training folds. The AP score is calculated on the held-out fold of each split. Individual scores across 5-fold cross-validation are shown as black dots above each bar representing the mean (n = 5 folds). (D) Evaluation of feature importance using the best-performing model, hyperparameter-optimized LightGBM. Shown are the prediction AP scores using training data from the indicated measurement(s). (E) Feature exclusion analysis to assess the contribution of individual features. Shown are changes in macro AP when the model was trained on all measurements except the indicated one(s). (D, E) Values are the mean ± SD over 10-fold cross-validation (n = 10 folds). In (E), positive values indicate a decrease in macro AP when the measurement is excluded. “Cell” refers to measurements across the entire cell.

As a demonstration, we compared 10 models: random forest, multilayer perceptron, extra trees, logistic regression, linear SVC, nearest neighbors, decision tree, ridge classifier, LightGBM, and gaussian naive-Bayes (Ke *et al*, 2017; Pedregosa *et al*, 2011) (Fig. 2A,B). The QuantCell framework readily accommodates additional models as appropriate. ML models were further optimized through hyperparameter tuning (Fig. 2A, Table EV2). This fine-tuning process is essential to optimize each model’s performance, ensuring the selection of the most effective model. During model selection and parameter tuning, principal component analysis (PCA) plots are generated to enable visual assessment for each cell type using the corresponding rule-based annotation markers (Appendix Fig. S3). These PCA plots facilitate model selection and parameter tuning by enabling visualization of the overlap between binary marker annotated cells and QuantCell-annotated cells in PCA space, as well as their separation from cells not assigned to that cell type.

While the optimal model selected by QuantCell may vary depending on the dataset, our analysis using bone marrow PhenoCycler data showed that the LightGBM model outperformed all other models tested after hyperparameter tuning (Figs 2A, B) while the random forest model performed best with default settings and required less runtime than LightGBM (Fig. EV2). LightGBM and random forest models are both based on decision trees. Their strong performance is consistent with the marker-based structure of the QuantCell classification task, which relies on combinations of positive and negative marker expression to define cell phenotypes. LightGBM is a sequential, gradient-based method which builds deep, asymmetric trees while random forest operates by averaging the predictions of multiple decision trees trained on subsets of the data (Breiman, 2001; Ke *et al*, 2017). Both the LightGBM and random forest models are widely used for classification tasks in many applications due to their versatility and efficacy (Boulesteix *et al*, 2012; Chen & Ishwaran, 2012).

### Impact of the data sources and statistical descriptors on prediction precision

During the ML model selection and optimization, the model was trained on all imaging measurements obtained from QuPath. This comprehensive quantitative dataset included data captured from multiple cellular compartments, such as the membrane, cytoplasm, nucleus, and whole-cell region. For each compartment, multiple statistical descriptors were collected, including the minimum, maximum, mean, median, and standard deviation of fluorescence intensity for each protein marker. Understanding the impact of data quantity, data source, and statistical descriptor on prediction precision could help identify data types with the highest and lowest predictive value, including those with detrimental effects that should be excluded to enhance QuantCell performance. This analysis was conducted using the hyperparameter-optimized LightGBM model, which demonstrated the highest macro AP score in our evaluation (Fig. 2A). Measurements of all markers were included when comparing cellular compartments and statistical descriptors. Cellular compartments were defined using the default StarDist-based (Schmidt *et al*, 2018) cell segmentation in QuPath.

Our analyses suggest that prediction performance depends on the amount of training data used (Fig. 2C). Comparing various data sources and statistical descriptors, our results suggest that the model performed the best when incorporating all cellular compartments and statistical measurements (Fig. 2D). Among the statistical measurements, the mean produced the best prediction performance, followed by the median, while the minimum performed markedly worse (Fig. 2D).

Comparing data from different cellular compartments, we found that the intensity measured across the entire cell generated the highest AP (Fig. 2D). Although most of the protein markers analyzed are localized to the cell surface, our results show that features derived from the cell membrane exhibited the lowest predictive performance (Fig. 2D). This is likely due to the dense cellular architecture of the BM, where closely packed cells can lead to overlapping membrane signals from adjacent cells. As a result, membrane measurements may not accurately represent protein expression levels in individual cells. In addition, our imaging data were acquired using a wide-field microscope rather than a confocal microscope. As a result, the nuclear region likely includes signals originating from the cytoplasm and membrane in the z-dimension.

Because some image analysis pipelines do not provide all types of measurements, we performed a feature exclusion analysis to assess how omitting specific types of imaging measurements affects model performance. Overall, the impact of excluding any single feature type, for all 25 protein markers, was negligible (Fig. 2E). Notably, removing features with high individual importance, such as the median or mean intensity of all cellular compartments, or all statistical measures across the whole-cell region (Fig. 2D), resulted in only minor reductions in the macro AP (Fig. 2E). The limited effect of feature exclusion may reflect overlapping predictive information among intensity descriptors and cellular compartments.

### Evaluation of QuantCell prediction performance

Using the hyperparameter-optimized LightGBM model and imaging data from all cellular compartments and statistical measurements, we next evaluated the predictive performance of QuantCell. We began by examining potential misclassifications using the reserved test dataset annotated by the binary marker rule-based method (Fig. 3A). While any classification strategy may contain errors, the rule-based annotations were used to train QuantCell and therefore served as the ground truth for this analysis.

**Figure 3.**
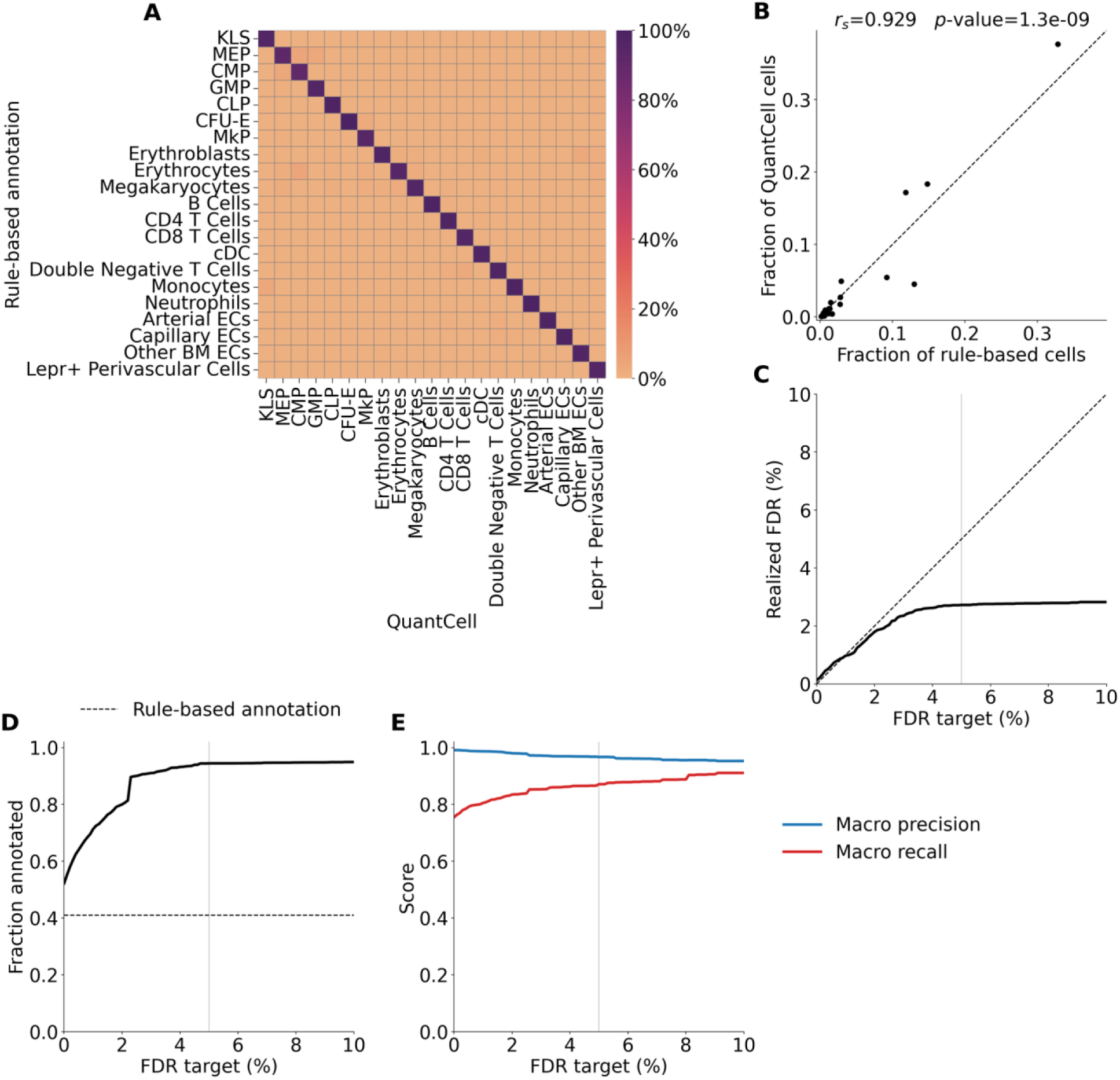
Evaluation of QuantCell prediction performance. (A) Column-normalized confusion matrix comparing rule-based cell annotation with QuantCell annotation. Values on the diagonal are equivalent to precision. n = 20,038 annotated test cells. (B) Comparison of cell type frequencies from rule-based and QuantCell annotations, calculated for each method as the fraction of annotated cells assigned to each cell type. Each dot represents one cell type (n = 21). Spearman’s rank correlation coefficient (rₛ) and two-sided *p*-value are shown (rₛ = 0.93, p = 1.3 × 10⁻⁹). The dashed line indicates equality between methods (*y* = *x*). (C) Targeted FDR used for thresholding versus pooled realized FDR across all cell types in the test set. The dashed line indicates equality (*y* = *x)*. (D) Targeted FDR versus the fraction of the full dataset annotated by QuantCell and the rule-based method. The dashed line represents the fraction annotated by the rule-based method alone. (E) Targeted FDR versus macro averaged precision and recall in the test set. (C–E) The light gray line represents the 5% FDR target used in this study. Values are from a single model evaluated on the held-out test set (n = 20,154 cells); no error bars.

Our results show that 97.3% of cells classified by QuantCell were consistent with annotations from the rule-based method, demonstrating high prediction accuracy across all cell types (Fig. 3A, Appendix Fig. S5). Only a small number of misclassifications were observed, primarily among closely related cell populations with overlapping marker-expression profiles (Table EV1). This number was measured using the fully held-out cell population in the test set that was not used for model training or threshold selection and included only cells with confident classifications under the 5% FDR threshold. Precision was also high across all cell types underscoring the model’s robustness even in unbalanced cell populations (Fig. 3A, Appendix Fig. S5) with a minimum precision of 0.91. The cell type frequencies predicted by QuantCell were consistent with those obtained from the rule-based annotation (Fig. 3B, Appendix Fig. S6). At a 5% FDR target, the realized FDR in the test set was 2.7% (Fig. 3C), and QuantCell increased the number of annotated cells from 41.0% to 94.4% (Fig. 3D), with macro precision of 0.97 and macro recall of 0.87 (Fig. 3E).

### Comparison of QuantCell with existing annotation frameworks

To evaluate QuantCell’s performance, we compared it with other cell annotation models developed for proteomics data, which are difficult to annotate due to limited marker availability (Table 1). Astir is a probabilistic model that uses deep recognition neural networks to annotate cell types based on user-defined markers for each cell type (Geuenich *et al*, 2021). AnnoSpat is a recently proposed semi-supervised approach that first clusters cells, then identifies a subset of confidently annotated cells using predefined positive and negative protein markers (Mongia *et al*, 2024). These annotated cells are used to train a neural network, which subsequently annotates the remaining cells. MAPS is another recently introduced model that uses a feedforward neural network trained on a ground truth dataset to perform cell type annotation (Shaban *et al*, 2024). CELESTA is a prior-based method which identifies high-confidence cells based on a cell type signature matrix and then uses spatial information to annotate neighboring cells (Zhang *et al*, 2022).

**Table 1.** Comparison of QuantCell with existing annotation frameworks.

| **Model** | **Method** | **Input data** | **Manual cell type annotation required** | **Marker combos** | **Uncertainty (allows unlabelled)** | **Macro precision** |
|---|---|---|---|---|---|---|
| **AnnoSpat** | Clustering | Expression-by-cell matrix & marker combos | No | Positive & negative | Yes | 0.304 |
| **Astir** | Clustering | Expression-by-cell matrix & marker combos | No | Positive | Yes | 0.352 |
| **CELESTA** | Iterative scoring with spatial neighbours | Expression-by-cell matrix, cell coordinates & marker combos | No | Positive & negative | Yes | 0.707 |
| **MAPS** | Feed-forward neural network | Expression-by-cell matrix & cell size | Yes | No | No | 0.759 |
| **QuantCell** | Variable classifiers | Expression-by-cell matrix & marker combos | No | Positive & negative | Yes | **0.967** |

We compared each model’s performance on the held-out test dataset using macro precision. Precision was selected as the primary benchmarking metric due to the importance of minimizing false-positive classifications, especially for rare cell types. Our analysis shows that QuantCell achieves higher precision than the other models evaluated (Fig. 4A, Fig. EV3), while also successfully annotating a substantial number of cells that cannot be classified by rule-based approaches (Fig. 3D). While other models exhibit a clear bias toward a few highly abundant cell populations with higher than 0.1 frequency (Fig. 4B), QuantCell maintains high precision across all cell types, including those with critical biological functions such as HSPCs (KLS and CLP) and their niche cells (e.g. Lepr^+^ perivascular cells; Ding *et al*, 2012; Zhou *et al*, 2014), where other models perform substantially worse (Fig. 4B). In addition, QuantCell operates around 7-fold faster than the second most accurate model, MAPS, in our dataset (Fig. 4C).

**Figure 4.**
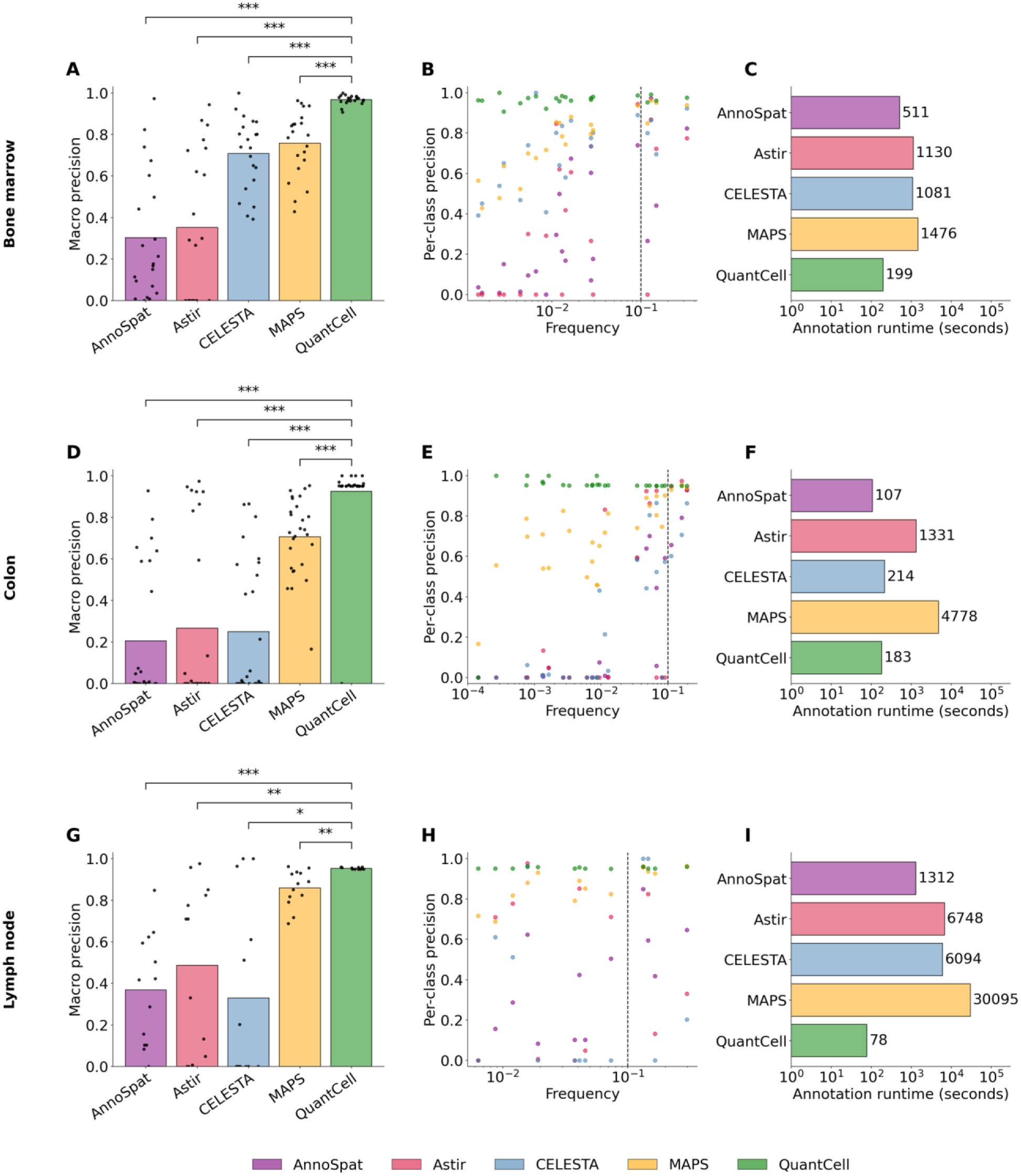
Benchmarking QuantCell against other spatial proteomics annotation methods. (A) Performance comparison using mouse bone marrow PhenoCycler data. Each dot represents a cell type. (B) QuantCell maintains high precision across cell types, whereas other methods show reduced precision for low-abundance cell types. Each dot represents a cell type. (C) Runtime required for model training and annotation of unknown cells. (D–F) Benchmarking model performance on an independent PhenoCycler colon dataset with previously established ground truth annotations (Schürch *et al*, 2020; Data ref: Schürch, 2020). (G–I) Benchmarking model performance on an independent multiplexed ion beam imaging (MIBI) lymph node datasets with previously established ground truth annotations (Shaban *et al*, 2024; Data ref: Shaban *et al*, 2023). (A, D, G) Bars show the macro average across cell types. Statistical significance was determined using a two-sided Wilcoxon signed-rank test with Bonferroni correction for four comparisons (*p < 0.05, **p < 0.01, ***p < 0.001). Exact Bonferroni-corrected p values: BM (n = 21 cell types): MAPS 3.8 × 10⁻⁶, CELESTA 7.6 × 10⁻⁶, Astir 3.8 × 10⁻⁶, AnnoSpat 3.8 × 10⁻⁶; colon (n = 27 cell types): MAPS 5.2 × 10⁻⁶, CELESTA 3.3 × 10⁻⁵, Astir 4.7 × 10⁻⁵, AnnoSpat 3.3 × 10⁻⁵; cHL1 (n = 13 cell types): MAPS 0.002, CELESTA 0.014, Astir 0.0029, AnnoSpat 0.00098. (C, F, I) Runtimes are from single runs; no error bars. (B, E, H) Dashed line marks a frequency of 0.1. QuantCell’s selected model was LightGBM for the BM and lymph node datasets, and random forest for the colon dataset.

We further benchmarked QuantCell against the other models using independent datasets generated from colon and lymph node tissues. The colon dataset is derived from primary human colorectal cancer samples and used PhenoCycler (previously known as Codex) for imaging (Schürch *et al*, 2020). The lymph node datasets were generated from human classical Hodgkin Lymphoma lymph node tissue and consist of two datasets imaged by mass spectrometry–based Multiplexed Ion Beam Imaging (MIBI), cHL1 and cHL2, and one imaged by PhenoCycler, cHL CODEX. All independent datasets included ground-truth annotations from the original studies (Schürch *et al*, 2020; Data ref: Schürch, 2020; Shaban *et al*, 2024; Data ref: Shaban *et al*, 2023). Our results demonstrate that QuantCell achieves higher precision, maintains robust performance across all cell types, and substantially reduces runtime across all independent datasets (Fig. 4, Fig. EV4) compared to MAPS, the next leading method. While other models exhibit a clear bias toward a few highly abundant cell populations with higher than 0.1 frequency, QuantCell delivers consistently high precision for functionally critical rare cell populations that can disproportionately influence disease progression and immune responses (Nguyen *et al*, 2018; Patel *et al*, 2024) (Fig. 4B,E,H, Tables EV3 and EV4).

Multilayer perceptron and MAPS are feed-forward neural networks, and both demonstrated relatively strong performance in our evaluation (Figs 2B, 4). However, MAPS relies on manually annotated cells for training, which is a major limitation (Shaban *et al*, 2024) (Table 1). Manual cell type annotation is labor-intensive for cell types defined by multiple markers and unfeasible for rare cell populations. In contrast, QuantCell automatically generates training datasets based on binary molecular marker profiles, eliminating the need for manual cell type annotation. Furthermore, QuantCell evaluates multiple ML models during model selection, enabling robust adaptation to diverse data characteristics.

The other three benchmarked methods, Astir, AnnoSpat, and CELESTA also have limitations in generating training datasets. AnnoSpat’s reliance on clustering compromises its performance in identifying rare cell types. While clustering is useful for pattern discovery, this unsupervised approach lacks the sensitivity of supervised learning, particularly in imbalanced datasets where rare cell types can be absorbed into larger, heterogeneous clusters (Weber & Robinson, 2016). Furthermore, clustering treats all markers as equally informative. Noise from uninformative markers can compromise the clustering and annotation accuracy. Astir is constrained by its use of only positive markers for annotation (Geuenich *et al*, 2021) (Table 1), assuming that marker expression is higher in the corresponding cell types than in all other cells. This assumption is not valid for markers shared by multiple cell types, which is common among stem and progenitor cells and diseased cells. CELESTA first annotates high-confidence “anchor cells” (Zhang *et al*, 2022) and then infers the identities of unannotated cells based on the cell types in close proximity. This reliance on spatial proximity may limit its performance in tissues with less spatial organization, such as BM, and for rare cell types that are spatially dispersed.

### Cell type specific prediction performance

A major strength of QuantCell is its strong performance across cell types with widely varying frequencies. We further evaluated QuantCell’s performance across individual cell types using the F1 score, the harmonic mean of precision and recall which summarizes predictive performance for each cell type. Our results suggested that the F1 score was significantly influenced by cell frequency and the number of protein markers used in rule-based annotation (Fig. 5A), likely due to their effects on the sample size for model training. In contrast, features such as cell size and cell circularity did not show direct correlation with F1 score (Fig. 5A), indicating the robustness and consistency of QuantCell’s performance across cell morphology.

**Figure 5.**
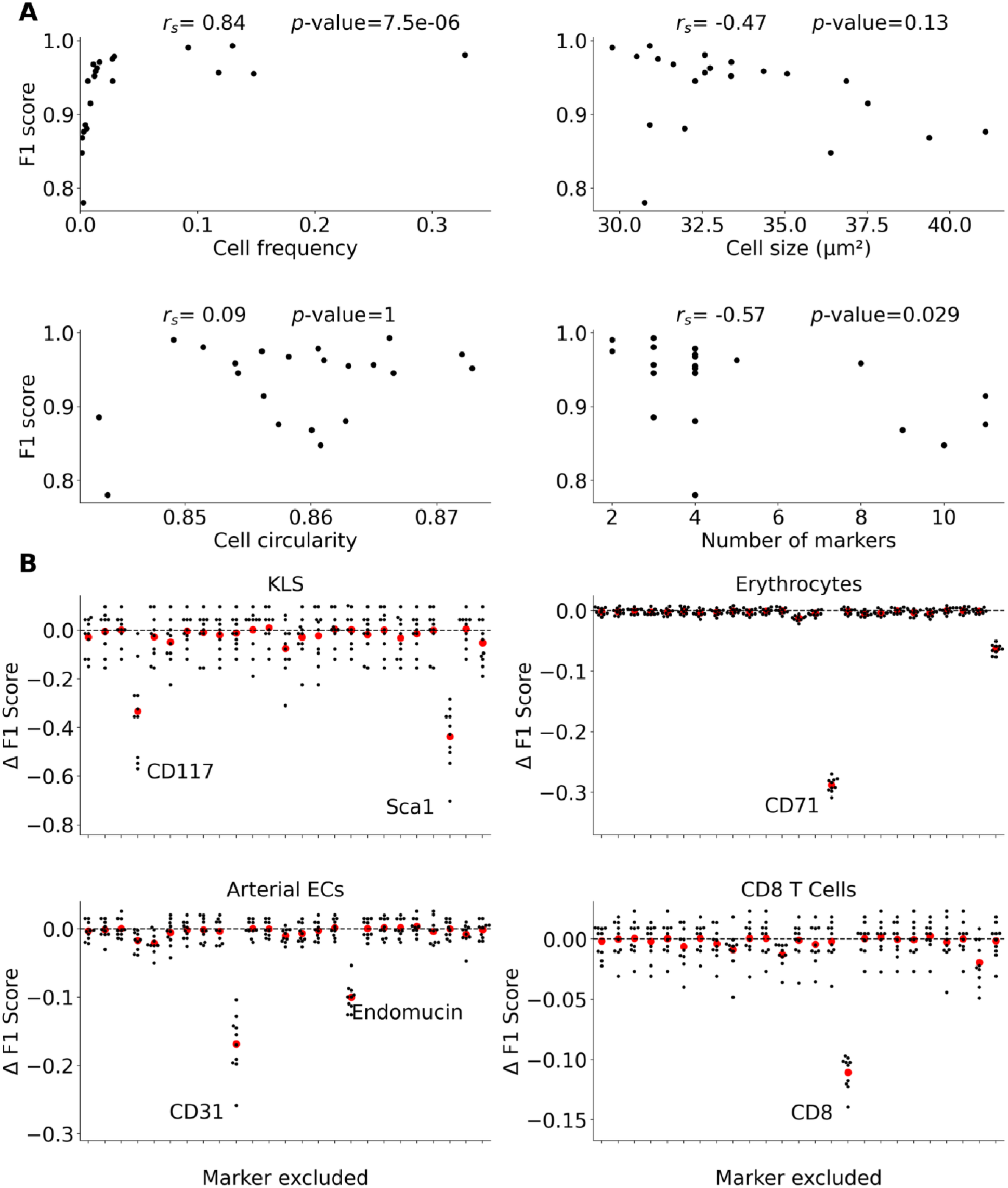
QuantCell prediction performance across cell types. (A) Comparison of F1 scores without FDR thresholding based on the cell type specific attributes: cell frequency, cell size, cell circularity, and the number of positive and negative markers used in rule-based annotation. Each dot represents one of 21 distinct cell types. Two-tailed Spearman’s rank correlation coefficients (rₛ) and corresponding *p*-values are shown with Bonferroni correction for four comparisons (cell frequency rₛ = 0.84, p = 7.5 × 10⁻⁶; cell size rₛ = −0.47, p = 0.13; cell circularity rₛ = 0.09, p = 1; number of markers rₛ = −0.57, p = 0.029). (B) Impact of individual protein markers on the prediction performance for representative cell types. Shown are changes in F1 score resulting from the exclusion of individual protein markers from the training and test datasets. Each dot represents the change in the F1 score relative to the baseline (no exclusion), calculated using 10-fold cross-validation. Red dots indicate the mean change across all 10-folds. Prominent markers are labeled.

To assess the contribution of individual protein markers to cell type classification, we systematically excluded one marker at a time from both the training and test datasets. This analysis revealed the markers that are indispensable for accurately annotating specific cell types (Fig. 5B, Appendix Fig. S7). As expected, these essential markers correspond to those used in rule-based annotation (Table EV1). However, not all markers used to define a given cell type have a strong effect. This discrepancy reflects how the model weighs informative features for each cell type and the redundancy among markers. For example, removing one lineage marker may have little effect on HSPCs because other lineage markers provide similar information. Together, these results revealed essential and redundant markers for cell type classification by QuantCell, which could help improve the design of marker panels.

## Discussion

In this study, we developed QuantCell, a ML based framework designed to improve cell annotation using qualitative and quantitative imaging data. We validated the performance of QuantCell using multiple complementary approaches. First, we assessed its accuracy on an independently held-out test dataset generated through binary marker rule-based annotation (Fig. 3A, C, E, Appendix Fig. S5). Second, all marker and cell annotations were independently reviewed by three field experts. Third, we evaluated QuantCell on four externally annotated datasets from published studies (Schürch *et al*, 2020; Shaban *et al*, 2024) generated using two multiplexed spatial imaging platforms, demonstrating robust, high-precision performance (Fig. 4D–I, Fig. EV4). Finally, PCA analyses showed that QuantCell-annotated cells tightly colocalize with binary marker annotated cells and are clearly separated from cells not assigned to the corresponding cell type (Appendix Fig. S3). Incorporating spatial context did not meaningfully improve cell annotation performance in our analysis of BM, a tissue with relatively disorganized architecture, lymph node, or colon, tissues with well-defined spatial organization (Fig. EV5). This may reflect the sparse spatial distribution of annotated cells, which limits the neighborhood information available to the model. Nevertheless, spatial features may provide additional predictive value in tissues with more pronounced spatial organization and in datasets containing a higher density of annotated reference cells.

QuantCell’s primary contribution is the integration of established analysis components into a practical cell-annotation workflow that offers the following advantages. First, QuantCell optimizes annotation performance across all cell types (Fig. 3A, Appendix Fig. S5). By avoiding performance bias toward abundant cell types (Fig. 4, Table EV4), it provides a more balanced and accurate classification. This is particularly important in biomedical datasets where rare cell types, such as stem cells or diseased cells, are often the primary focus (Nguyen *et al*, 2018; Patel *et al*, 2024) but are poorly detected by existing methods (Fig. 4B,E,H). Second, QuantCell performs annotation based on a user-defined FDR, enabling stringent control over classification confidence (Fig. 3C–E). It reduces the risk of misclassification by leaving uncertain cells unannotated, which improves the reliability of biological interpretation. Third, QuantCell does not require manual cell type annotation, which is labor-intensive for cell types defined by multiple markers and unfeasible for rare cell populations. Instead, it uses both positive and negative markers to guide training dataset generation, enabling evaluation of each marker’s contribution (Fig. 5B, Appendix Fig. S7). QuantCell is designed for image-specific model training and cell classification. Because marker-intensity distributions and other technical characteristics may vary across images and imaging runs, applying a trained model to an independent image may require image-specific retraining or data harmonization.

As a supervised approach, QuantCell is limited to cell populations specified in the annotation rules and is not designed for the de novo discovery of unannotated populations. Its performance also depends on the quantity and quality of the available training data. It may underperform on datasets with sparse or low-dimensional features (Fig. 2C, D) or for cell types with limited or low-quality training data (Fig. 5A). To improve the quality of the training and test datasets, one potential approach is to filter out outlier cells annotated by the rule-based method. Because the rule-based method assigns cell identities based on predefined marker combinations (Table EV1), cells of the same type are expected to cluster in the corresponding marker space (Appendix Fig. S3). By comparing the density of cells assigned to a given type with the densities of other types in this space, it is possible to estimate the probability that each cell truly belongs to its assigned identity. The cell distributions can be visually inspected in PCA plots to determine whether low-probability cells should be removed. Implementing such procedures in the future could further improve model predictions.

Despite this limitation, QuantCell is able to increase the number of annotated cells from 41.0% to 94.4% at a 5% FDR, achieving 97.3% overall accuracy based on the test dataset from rule-based annotation (Fig. 3A, D). Our benchmarking analysis shows that QuantCell outperforms existing platforms in prediction precision, particularly for low-frequency cell types (<0.1), including those with important biological functions such as stem cells and diseased cells (Fig. 4). It has the potential to be compatible with a wide range of spatial omics platforms, including two- and three-dimensional imaging, RNA-based technologies such as seqFISH (Lubeck *et al*, 2014) and MERFISH (Chen *et al*, 2015), and sequencing-based approaches like Visium (Ståhl *et al*, 2016), Slide-seq (Rodriques *et al*, 2019), and Stereo-seq (Chen *et al*, 2022), although the sparse nature of the data may present challenges. The framework is also adaptable to project-specific priorities. For example, QuantCell is configured by default to optimize performance equally across all cell types, but users can customize weighting schemes through the chosen classifier to prioritize specific cell populations of interest. Users can also incorporate their preferred supervised ML models in QuantCell for model selection (Fig. 1F). Future studies could further expand QuantCell’s capabilities by incorporating transfer learning to enhance performance in data-sparse settings. In addition, incorporating domain adaptation (Bae *et al*, 2022; Pan & Yang, 2010) techniques may enable QuantCell to generalize across tissue types, experimental platforms, or imaging modalities with minimal retraining.

In summary, QuantCell provides a configurable workflow that integrates established supervised ML methods to improve cell annotation. By integrating qualitative marker definitions with quantitative imaging features and applying a user-defined FDR to control classification confidence, QuantCell substantially increases the number of annotated cells while maintaining high accuracy. It outperforms existing annotation methods in prediction precision, minimizes performance bias toward abundant cell types, and enables robust detection of rare cell populations. Unlike many existing methods, QuantCell does not require manual cell type annotation, which can be labor-intensive for multi-marker cell types and impractical for rare cell populations. QuantCell is compatible with a wide range of spatial omics platforms and can be tailored to project-specific priorities. Its flexibility and precision enable accurate characterization of complex cellular phenotypes, including those with highly imbalanced cell populations. We have provided video tutorials to guide researchers in using QuantCell (links in the GitHub repository, see Data availability). QuantCell accepts appropriately formatted marker measurements generated using QuPath or other image-analysis tools. Because the best-performing ML models identified in our evaluation may not be optimal for other datasets or classification tasks, QuantCell allows users to compare candidate models and select the one that performs best for their specific dataset and research objective.

## Methods

### Reagents and Tools Table

The Reagents and Tools Table is provided as a separate file.

### Methods and Protocols

#### Mice

C57BL/6J mice (Jackson Laboratory stock #000664) were maintained in the Research Animal Facility at the University of Southern California (USC) under a 12:12 hour light-dark cycle, with controlled room temperature (22 ± 1 °C) and humidity (55 ± 10%). All animal procedures were approved by the USC Institutional Animal Care and Use Committee (protocols 21461 and 21684) and performed in accordance with the Guide for the Care and Use of Laboratory Animals (National Research Council, 2011).

#### PhenoCycler antibodies

The following primary antibodies were obtained from Akoya Biosciences: RX025-AF488 anti-CD11b (M1/70), RX030-Cy5 anti-CD11c (N418), RX020-ATTO550 anti-CD19 (6D5), RX021-Cy5 anti-CD3 (17A2), RX002-ATTO550 anti-CD31 (MEC13.3), RX026-ATTO550 anti-CD4 (RM4-5), RX007-AF488 anti-CD45 (30-F11), RX010-AF488 anti-CD45R (RA3-6B2), RX027-Cy5 anti-CD71 (RI7217), RX029-ATTO550 anti-CD8a (53-6.7), RX024-Cy5 anti-Ly6g (1A8), RX014-ATTO550 anti-MHCII (M5/114.15), RX003-Cy5 anti-TCRαβ (H57-597), and RX004-AF488 anti-TER119 (TER-119). The following primary antibodies were obtained from ThermoFisher Scientific: BX035-ATTO550 anti-CD105 (209701), BX050-AF647 anti-CD115 (AFS98), BX045-Cy5 anti-CD117 (2B8), RX013-AF488 anti-CD34 (RAM34), BX037-ATTO550 anti-CD41 (MWReg30), BX040-AF488 anti-endomucin (polyclonal), BX036-Cy5 anti-F4/80 (CI:A3-1), BX041-ATTO550 anti-CD127 (A7R34), and BX042-Cy5 anti-Sca1 (D7). The following primary antibody was obtained from Leinco Technologies: BX028-AF488 anti-CD32/CD16 (2.4G2). The following primary antibody was obtained from R&D Systems: BX049-ATTO550 anti-LEPr (polyclonal). All primary antibodies obtained from vendors other than Akoya Biosciences were conjugated to Akoya barcodes by Akoya Biosciences.

#### Antibody validation and quality control

All antibodies were conjugated and validated by Akoya Biosciences. Successful conjugation was verified by sodium dodecyl sulfate–polyacrylamide gel electrophoresis (SDS–PAGE), confirming that each conjugated antibody migrated at the expected molecular weight. After verification of conjugation, Akoya staff stained our tissue sections with the antibodies and evaluated antibody performance using a standardized scoring system: –1 (non-specific binding), 0 (no signal/no positive cells), 1 (weak signal), 2 (good staining), and 3 (saturated). Antibody clones and/or dilutions were adjusted based on these evaluations. A blank imaging cycle was used as a negative control. All antibodies included in the final panel received a rating of 2. Akoya personnel had no prior knowledge of the sample identities.

#### Bone marrow preparation for imaging

BM was carefully flushed from the diaphysis of femurs and tibias using a syringe filled with DPBS with 2% FBS, ensuring gentle handling to obtain the whole, intact BM plugs. The collected BM was immediately fresh-frozen in tissue freezing medium (Electron Microscopy Sciences, Cat. #72592) for downstream applications.

#### Bone marrow imaging

Glass coverslips, 22 mm x 22 mm, were pre-treated with poly-L-lysine (Santa Cruz Biotechnology, Cat. #sc-286689) to enhance tissue adherence. The fresh-frozen BM samples were cryosectioned at a thickness of 5 µm and mounted onto the prepared coverslips. Six BM section samples from six mice were imaged and processed, yielding a total of 245,719 cells identified through QuPath segmentation. Cells from all six samples were pooled for downstream analyses. Antibody staining and imaging were performed by Akoya Biosciences personnel using Akoya’s established protocols.

#### Cell segmentation

Nuclei-based cell segmentation was performed based on DAPI stains in QuPath (Bankhead *et al*, 2017) (version 0.3.0) using the StarDist (Schmidt *et al*, 2018) (version 0.3.0) algorithm, using the settings provided in our GitHub (see Data availability). For nuclei-based segmentation, QuPath automatically infers cytoplasm and membrane compartments by expanding nuclear boundaries up to 5 µm constrained to 1.5x the nucleus size. Membrane-based segmentation was performed with Cellpose-SAM (preprint: Pachitariu *et al*, 2025) (version 4.0.6) using a two-channel image of a composite of F4/80, CD45, Endomucin, Sca1, and CD71, along with DAPI.

To recover cellular compartments using Cellpose, segmentation was run independently on the composite (whole cell) and the DAPI channel (nucleus). The cytoplasmic compartment was defined by subtracting the nuclear area from the whole-cell area. For comparing the membrane-based and nuclei-based approaches, Cellpose cells with exactly one StarDist nuclear centroid were retained (194,149 of 272,315) and the rule-based annotation labels were matched from the StarDist set to the Cellpose set. QuantCell achieved comparable precision with either membrane-based or nuclei-based cell segmentation (Fig. EV1). Nuclei-based segmentation was used for all other analyses.

#### Binary marker rule-based annotation and QuPath settings

Cell type labels were assigned by matching each cell’s binary marker vector to the predefined marker combinations in Table EV1. This process is rule-based and deterministic, not model-based. Qualitative marker calling was performed in QuPath (Bankhead *et al*, 2017) (version 0.3.0). For each marker, approximately 20-70 positive and 20-70 negative cells from across the tissue section were manually annotated. A cell was considered positive for a marker when its signal clearly exceeded the background signal and exhibited the expected staining pattern for that marker. Signal adjustment and staining patterns were used to minimize the effects of non-specific binding and background signals. Then, QuPath’s built-in object classifier propagated these binary labels to all cells in the section. Propagated calls were reviewed, and if errors were identified, additional positive and negative training cells were added, and the classifier was rerun until the resulting cell annotations passed manual assessment. Mutually exclusive and co-expressing marker combinations were also used for further verification.

To reduce potential cross-contamination from neighboring cells (Bai *et al*, 2021), our annotation strategy incorporated the spatial distribution of marker signal within each cell. During manual binary marker assignment in QuPath, cells were classified as negative when the apparent signal was confined to regions adjacent to neighboring positive cells. In the QuantCell annotation, spatial distribution for each marker was quantified using multiple statistical metrics, including the standard deviation of signal intensity within each cellular compartment.

The resulting marker assignments were independently verified by three field experts, including a board-certified pathologist, the director of our imaging core facility, and an experienced tissue biologist, who qualitatively assessed whether the assigned cell type labels were consistent with the expected marker-expression patterns. After verification, the protein-expression–by–cell matrix and the binary positive/negative marker calling were exported from QuPath. The binary annotations were then used for the rule-based cell-type annotation. Detailed information on the procedure used to generate the protein-expression-by-cell matrix is available in our GitHub repository (see Data availability).

Using the binary positive/negative marker calls generated by QuPath, QuantCell constructed the training and test datasets by annotating cells that exactly matched predefined marker combinations (Table EV1). Cells with ambiguous profiles, such as those matching more than one combination or none at all, were not assigned a cell type and were labeled as “unannotated”. To further verify cell annotation accuracy, single-cell annotations generated by both the rule-based approach and by QuantCell were independently assessed by three experts as mentioned above (Appendix Fig. S4). Expert assessment is not required for QuantCell-based annotation, but was used here for independent validation of QuantCell’s performance. Note: QuPath is not required to run QuantCell and is not part of the QuantCell software. QuantCell also accepts appropriately formatted cell marker quantification data generated using other tools, provided that binarized marker data or a subset of annotated cells is available.

#### Data preprocessing

Cells that were successfully annotated by the binary marker rule-based method were split into an 80/20 train–test dataset using train_test_split() from scikit-learn (version 1.5.0) (Pedregosa *et al*, 2011), with stratification based on class distribution and a fixed random seed of 42. Features were normalized to have a mean of 0 and a standard deviation of 1 using StandardScaler() from scikit-learn. The scaler was fitted to the marker distributions in the training dataset and then applied to both the test dataset and unannotated cells without refitting, to prevent data leakage. Cell type labels were encoded using LabelEncoder() from scikit-learn after excluding any cells that were originally unannotated.

#### Model selection

For the BM dataset, initial model performance was evaluated across ten different base classifiers with default parameters, using the training dataset and 10-fold cross-validation. All ten models were further optimized for macro AP using HalvingGridSearchCV from scikit-learn (Jamieson & Talwalkar, 2016; Pedregosa *et al*, 2011), with 10-fold cross-validation on the training dataset. Search parameters are listed in Table EV2. For the external datasets, each tuned model was tested using the hyperparameters derived from BM and the best model was selected by macro AP. The tuned LightGBM (version 4.6.0; Ke *et al*, 2017) model had the highest macro AP for every dataset but random forest was used for colon because it assigned all but one cell type at 5% FDR compared to three cell types left unannotated by LightGBM.

#### Training size analysis

To evaluate the dependence of model performance on training data availability, we performed a training size analysis using 5-fold cross-validation. 5-fold cross-validation was used instead of 10-fold to match the analysis strategy used in Figure 4A of the MAPS manuscript (Shaban *et al*, 2024). Within each fold, the validation set was held constant to ensure paired comparisons, while the training set was randomly subsampled at varying proportions (5%, 10%, 25%, 50%, 75%, and 100%). Consequently, the 100% condition represents the full set of remaining 4-folds. All rule-based annotated cells, including the test dataset, were used for this analysis.

#### Feature analysis

Performance of the model was evaluated by systematically varying the inclusion or exclusion of specific measures or markers across 10-fold cross-validation on the training set. Each fold was treated as an individual data point. For the cell region and statistical analyses (Fig. 2D, E), performance was scored using macro AP. For marker analyses (Fig. 5B, Appendix Fig. S7), all features of each marker were excluded one at a time. Cell type F1 scores were calculated without applying a FDR threshold and were compared to an all-marker baseline. Markers were labeled when the mean change in F1 was at most -0.02 and the change was at least 25% of the largest decrease for that cell type. This criterion was arbitrarily chosen to highlight prominent markers.

For the cell type comparisons in Fig. 5A, cell frequency is calculated as the fraction of cells annotated using the rule-based method that belong to each cell type. Cell size and circularity are the mean whole-cell area and circularity for all cells of a cell type. The number of markers are the total number of positive and negative markers used in a marker combination for a cell type. Two-sided Spearman’s rank correlation test was performed for each panel across the 21 cell types. P-values were Bonferroni-corrected for the four features tested.

#### Model thresholding

In typical use, class-specific probability thresholds are determined from the empirical precision-recall curve at a user-specified FDR target using a 20% held-out test set, and then applied to the unannotated cell population. This procedure does not require classifier output probabilities to be well calibrated. The probability threshold is set per class to the lowest value at which interpolated precision is at or above 1 - FDR. If no class achieves this threshold, the class is left unannotated. Each cell is annotated as the most probable cell type if that probability was at or above the inferred threshold and left unannotated otherwise.

For the performance evaluation in Fig. 3A, C and E, threshold selection and accuracy assessment were performed on separate datasets. The original 20% test set was completely excluded from model development. The remaining 80% of labeled cells was further split 80/20 into a training set (used for model fitting) and a validation set (used for threshold selection). The selected thresholds were then applied to the held-out test set, whose labels were revealed only for the final accuracy comparison. For thresholded analyses in this study, the default FDR target was set to 5%. Thresholding was not applied in any analysis involving the AP score, as AP is calculated by evaluating performance across all possible thresholds.

#### Benchmarking QuantCell against existing models

For all benchmarking analyses, annotations from the binary marker rule-based method were used as the ground truth. For Astir (version 0.1.2; Geuenich *et al*, 2021), AnnoSpat (version 1.0.0; Mongia *et al*, 2024), MAPS (version 1.0.0; Shaban *et al*, 2024), and CELESTA (R 4.4.2; https://github.com/plevritis-lab/CELESTA; Zhang *et al*, 2022), mean expression of each marker across the entire cell was used as input, following the pipeline described in the corresponding papers. MAPS was also provided the cell area (µm^2^) as directed.

CELESTA (R 4.4.2) was run on all cells separately for each tissue section. Cell centroid coordinates and mean whole-cell marker expression were used as inputs along with a prior signature matrix built from the same marker combinations as QuantCell. Default filtering thresholds were used.

Astir and AnnoSpat were applied to the entire rule-based annotated dataset. No preprocessing was performed. The models were run with default parameters. The same marker combinations used in QuantCell were applied, except negative markers were excluded for Astir, as they are not supported by this model.

MAPS was trained and tested using the same 80/20 train–test split as QuantCell. Labels were encoded as in the QuantCell workflow prior to training. No preprocessing was applied, and default parameters were used, except for “min_epochs” which was reduced to 5 and “patience” which was reduced to 25 to avoid unnecessary computation.

QuantCell was trained and tested using the same 80/20 train–test split as MAPS. Performance benchmarking was conducted with thresholding at 5% FDR.

All annotation methods were evaluated on the same held-out test set (20,154 cells). Cells that were unannotated by a method were counted as “missed” for recall and excluded from the precision calculation.

#### Benchmarking on external datasets

We benchmarked QuantCell against each annotation method using four external datasets from two tissue types. The colon dataset used CODEX (PhenoCycler) to profile colorectal cancer samples (Schürch *et al*, 2020; Data ref: Schürch, 2020). Lymph node data consisted of three classical Hodgkin lymphoma (cHL) datasets from the original MAPS study (Shaban *et al*, 2024; Data ref: Shaban *et al*, 2023): two generated using multiplexed ion beam imaging (MIBI) and one using CODEX (PhenoCycler). In these datasets, ground truth annotations were established through cluster-based annotation followed by manual curation (Schürch *et al*, 2020; Shaban *et al*, 2024). Marker combinations for Astir, AnnoSpat, and CELESTA were based on the original studies (Schürch *et al*, 2020; Shaban *et al*, 2024).

QuantCell was assessed directly against the provided ground truth annotations. To ensure a fair comparison, QuantCell was configured using the same 80/20 train–test split as MAPS, and performance was evaluated at a 5% FDR threshold with class thresholds determined on the held-out test cells, as in typical use (see Model thresholding). MAPS was rerun in the same computational environment rather than using the performance metrics reported in the original publication (see Computational environment). QuantCell used LightGBM for all three cHL datasets and random forest for the colon dataset. Reported runtime includes both model training and prediction. For Fig. 4 and Fig. EV4, statistical significance was determined using a two-sided Wilcoxon signed-rank test comparing QuantCell with each other method. A Bonferroni correction was applied for four comparisons.

#### Spatial feature analysis

To test whether adding spatial features would improve QuantCell performance (Fig. EV5), cell-centered neighborhood features were added to the standard feature set. For every cell, the nearest k cells (k=10 or 20) were detected using a k-d tree (SciPy 1.16.3 cKDTree; Virtanen *et al*, 2020) within each section (BM), region (colon), or stitched field (cHL1). Three sets of features were computed: (i) neighbor-mean intensities, the mean intensity across the whole cell for each marker, (ii) rule-based composition, the fraction of k neighbors that were a cell type for each cell type annotated by the rule-based method (for BM only), and (iii) local density, the mean distance to the k neighbors as well as the number of cells within 30 µm. “All families” represents that every spatial feature set was included. Performance was evaluated without FDR thresholding after fitting to the training dataset and evaluating on the test dataset.

#### FDR sweep

For the FDR analyses in Fig. 3C–E, thresholds were selected in steps of 0.1% from 0% to 10%. Realized FDR was calculated as the fraction of QuantCell-annotated cells in the test set whose label differed from the rule-based method.

#### Principal component analysis

For each cell type, cells were separated into three groups: cells annotated as that type by the rule-based method, cells annotated as that type by QuantCell only, and cells not assigned that type by either method. From each group, 500 cells were sampled with replacement and a fixed seed. Principal component analysis was performed using unscaled intensity measurements across all cell regions and statistics for only the markers which are in that cell type’s marker combinations. Only the first two components are shown.

#### Computational environment

All benchmarking tests and analyses were performed on a dedicated server with Ubuntu 20.04.6 LTS, 2 x Intel(R) Xeon(R) Gold 6136 CPU @ 3.00GHz (24 physical cores, 48 threads total, 49.5 MB L3 cache), 494.7 GB memory. All code is in Python, except for CELESTA (R 4.4.2) and a Bash script used to run AnnoSpat.

#### Statistics and reproducibility

Statistical tests used in data analysis are detailed in the relevant figure legends and “Methods” sections. No statistical method was used to predetermine sample size. Expression data from markers with poor staining quality or those not included in any marker combinations (Table EV1) were excluded from analysis. Where applicable, 10-fold cross-validation was employed for model selection. A fixed random seed was applied during data splitting to ensure reproducibility across sessions.

Investigators were not blinded during sample collection and data analysis.

## Data availability

The datasets and computer code produced in this study are available in the following databases:

- PhenoCycler image, QuPath exports and QuantCell input and output tables: Zenodo 10.5281/zenodo.23672666 (https://doi.org/10.5281/zenodo.23672666) (Data ref: Boohar et al, 2026)
- QuantCell code and the analysis scripts for this paper (“paper” directory): GitHub ronglulab/QuantCell (https://github.com/ronglulab/QuantCell)

The colon dataset was obtained from Mendeley Data (Data ref: Schürch, 2020) and the three cHL datasets from Zenodo (Data ref: Shaban et al, 2023).

## Author contributions

W.B. and R.L. conceived and designed the project. B.W. designed the antibody marker panel and cell type marker combinations, conducted all wet lab experiments including BM sectioning. B.W. also performed cell segmentation and marker calling in QuPath. W.B. analyzed the data and wrote all project-related code. B.W., Z.T., A.N., and R.L. assisted with data analysis. W.B. and R.L. wrote the manuscript. All authors reviewed and approved the final manuscript.

## Acknowledgements

We thank Akoya Biosciences for their assistance in establishing the antibody panel and acquiring imaging data, and Drs. G. Fudenberg, A. MacLean as well as the members of the Lu laboratory for their valuable discussions and insights. We also thank Drs. S.W. Ruffins and I. Hajjali for assistance with validating marker and cell annotations.

This study is supported by R35HL150826, R01DK143671, and R01AG080982. W.B. was supported by USC Provost’s Fellowship, QCB Google Alumni Award in Quantitative Biology, and the John Petruska Memorial Award.

---

**Figure EV1.**
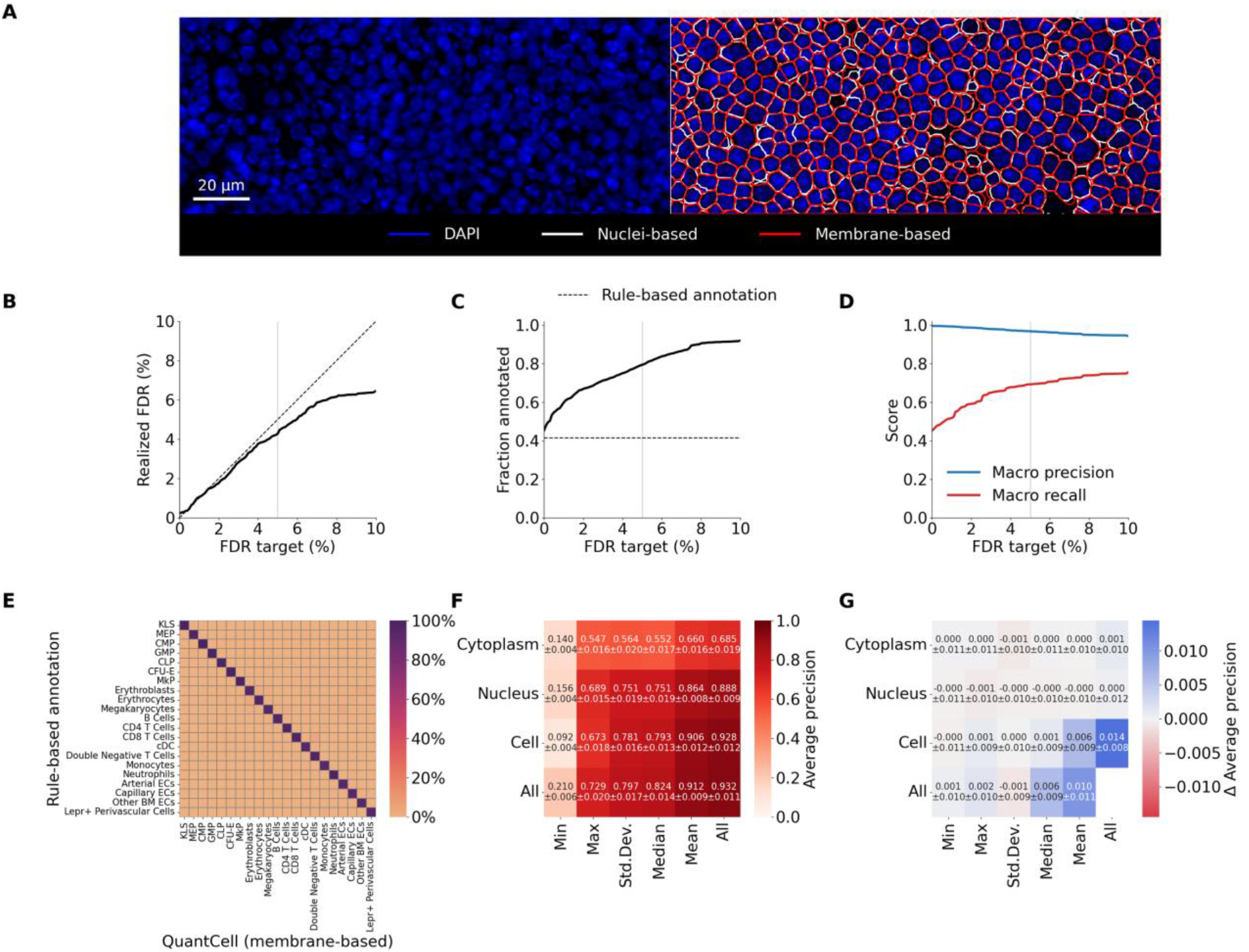
QuantCell performs comparably using nuclei-based and membrane-based cell segmentation. (A) Overlay of nuclei-based and membrane-based segmentation results. Image acquired at 20X magnification. Scale bar, 20 µm. (B–G) Analyses using membrane-based cell segmentation. (B) Targeted FDR used for thresholding versus pooled realized FDR across all cell types in the test set. Dashed line indicates equivalence. (C) Targeted FDR versus the fraction of the full dataset annotated by QuantCell and the rule-based method. The dashed line represents the fraction annotated by the rule-based method alone. (D) Targeted FDR versus macro averaged precision and recall in the test set. (E) Column-normalized confusion matrix comparing rule-based cell annotation with QuantCell annotation. Values on the diagonal are equivalent to precision. n = 14,423 annotated test cells. (F) Evaluation of feature importance using the best-performing model, hyperparameter-optimized LightGBM. Shown are the prediction AP scores using training data from the indicated measurement(s). (G) Feature exclusion analysis to assess the contribution of individual features. Shown are changes in macro AP when the model was trained on all measurements except the indicated one(s). (B–D) Light gray line represents the 5% FDR target used for this paper. Values are from a single model evaluated on the held-out test set (n = 16,108 cells); no error bars. (F, G) Values are the mean ± SD over 10-fold cross-validation (n = 10 folds). In (G), positive values indicate a decrease in macro AP when the measurement is excluded. “Cell” refers to measurements across the entire cell.

**Figure EV2.**
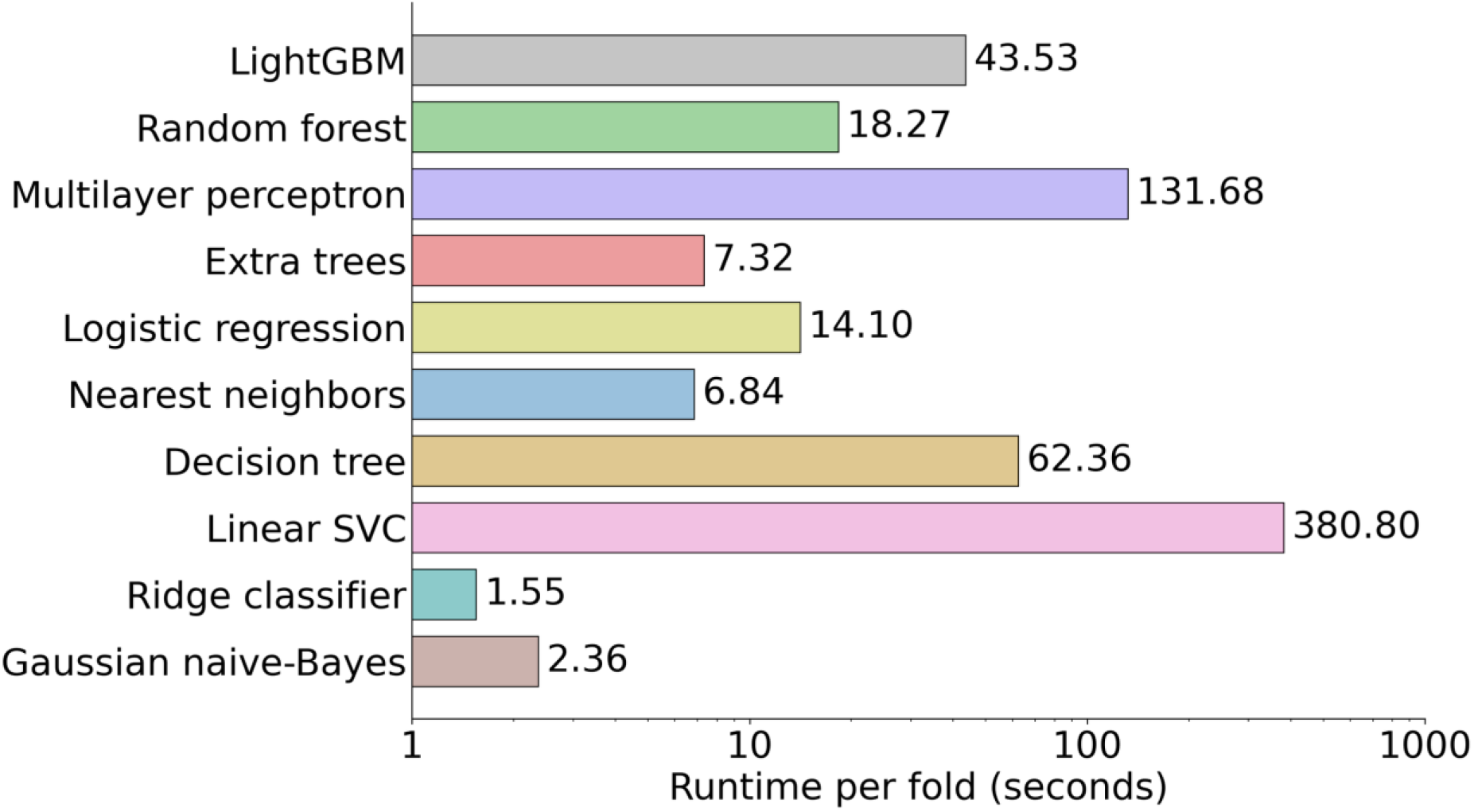
Runtime comparison of base classifier models. Mean runtime per fold for base classification models on the data used in this paper. Bars show the mean per-fold runtime over 10-fold cross-validation (n = 10 folds); no error bars.

**Figure EV3.**
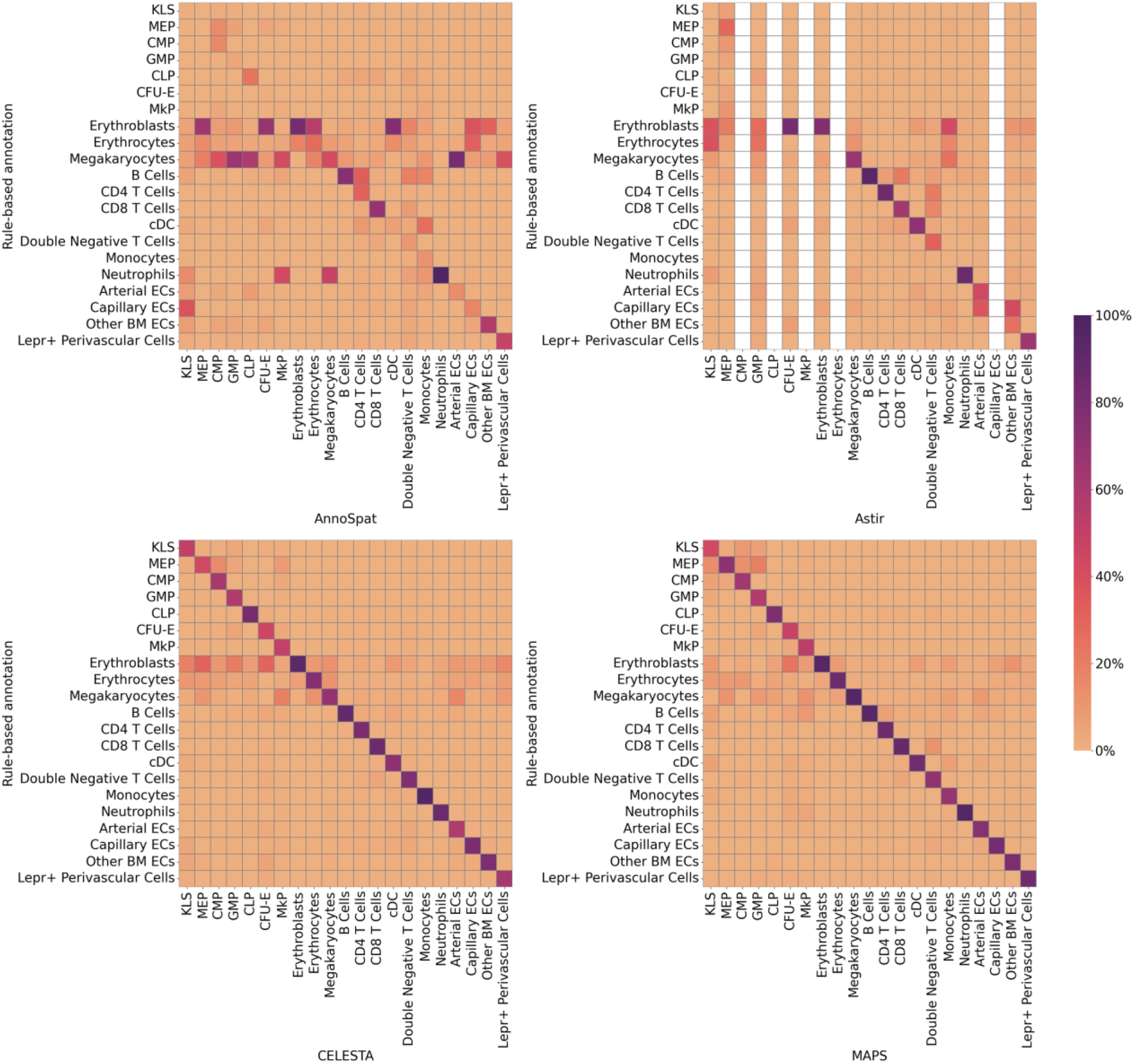
Confusion matrices for predictions generated by AnnoSpat, Astir, CELESTA, and MAPS in the BM dataset. Blank columns represent that the model did not annotate any cells with that cell type. Matrices are column-normalized. AnnoSpat, Astir and CELESTA were evaluated on all rule-based annotated cells (n = 100,769); MAPS was evaluated on the held-out test cells (n = 20,154).

**Figure EV4.**
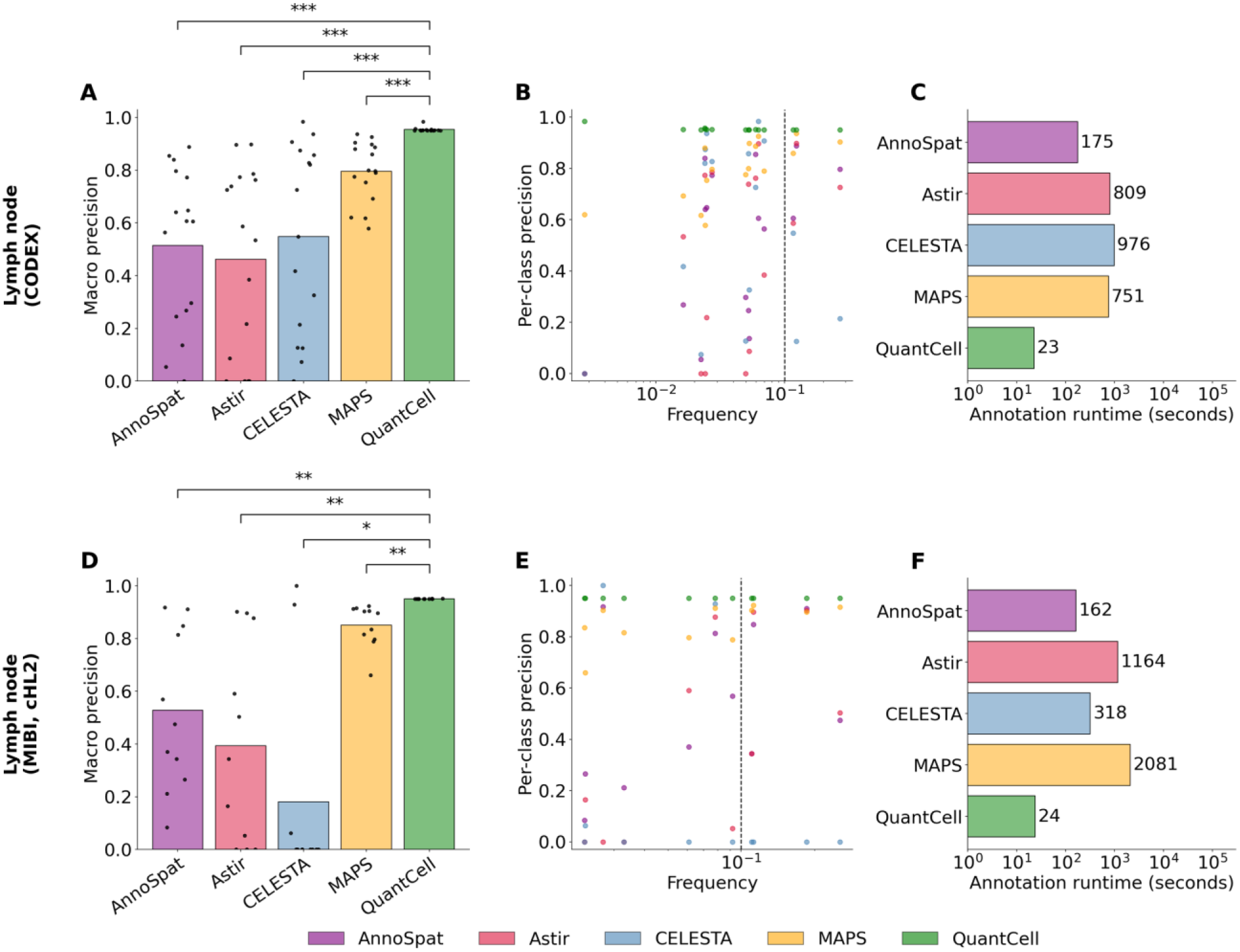
Benchmarking on additional lymph node datasets. (A–F) Performance comparison across two additional lymph node datasets (Shaban *et al*, 2024; Data ref: Shaban *et al*, 2023). Methodology is the same as Fig. 4. In (A, D), bars show the macro average across cell types. Statistical significance was determined as in Fig. 4 (two-sided Wilcoxon signed-rank test, Bonferroni correction for four comparisons); exact corrected p values: cHL CODEX (n = 16 cell types): MAPS 0.00012, CELESTA 0.00037, Astir 0.00012, AnnoSpat 0.00012; cHL2 (n = 11 cell types): MAPS 0.0039, CELESTA 0.012, Astir 0.0039, AnnoSpat 0.0039.

**Figure EV5.**
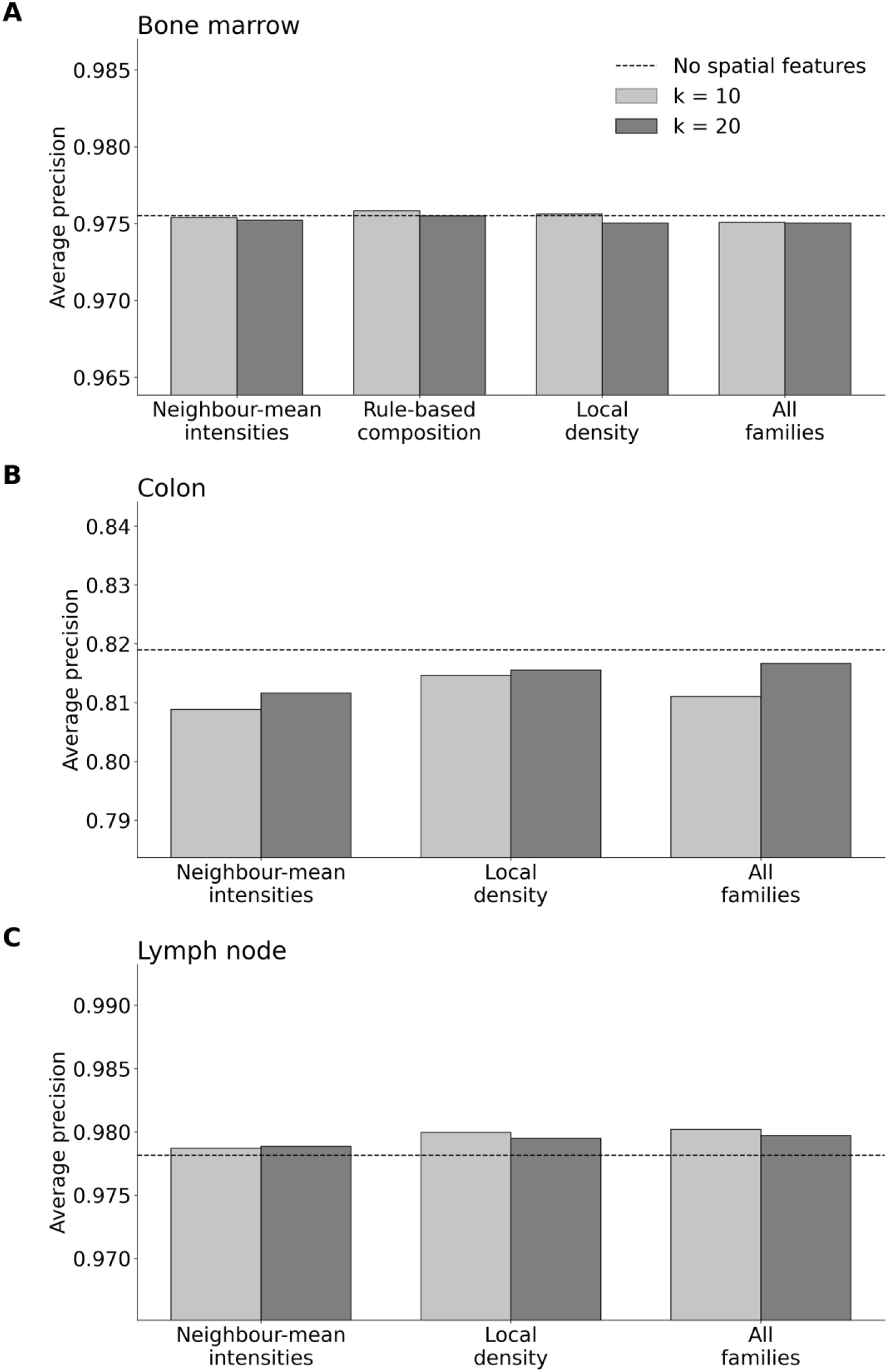
Impact of spatial feature data on model performance. Macro AP of the QuantCell model with varying spatial features added in addition to the standard set for (A) bone marrow, (B) colon, and (C) lymph node (cHL1) datasets. Dashed line represents standard model performance. LightGBM was used for all three datasets. Neighbor-mean intensities: mean intensity for each marker in the k neighbors surrounding a cell. Rule-based composition: fraction of k neighbors that are each cell type. Local density: mean distance to k neighbors and number of cells within 30 micrometers. Colon and lymph node datasets have their own ground truth labels and do not use rule-based annotation so rule-based composition is excluded. Each bar is a single model fit evaluated on the held-out test set (bone marrow n = 20,154, colon n = 48,901, cHL1 n = 307,687 cells); no error bars.

**Table EV1.** Marker combinations for rule-based annotation. Marker combinations for rule-based annotation. *the listed marker combination also detects a subset of monocyte-derived macrophages.

| cell type | label in QuantCell | positive markers | negative markers | PMIDs | reference URLs |
|---|---|---|---|---|---|
| Arterial Endothelial Cells (Arterial ECs) | Arterial ECs | CD31 | CD117, CD11b, Endomucin, TCR | 31871321, 16166583 | https://pmc.ncbi.nlm.nih.gov/articles/PMC7610809/; https://ashpublications.org/blood/article/107/2/535/131454/Platelet-PECAM-1-inhibits-thrombus-formation-in |
| B Cells | B Cells | B220, CD19 |  | 39266537 | https://www.nature.com/articles/s41467-024-52224-6 |
| Capillary Endothelial Cells (Capillary ECs) | Capillary ECs | Endomucin | CD117, CD11b, TCR | 24646994, 27074508, 36805714 | https://pmc.ncbi.nlm.nih.gov/articles/PMC4943525/; https://pmc.ncbi.nlm.nih.gov/articles/PMC5035541/; https://pmc.ncbi.nlm.nih.gov/articles/PMC9942337/ |
| CD4 T Cells | CD4 T Cells | CD3, CD4, TCR | CD8α | 20556631, 34524664 | https://pmc.ncbi.nlm.nih.gov/articles/PMC11115818/; https://pmc.ncbi.nlm.nih.gov/articles/PMC10802914/ |
| CD8 T Cells | CD8 T Cells | CD3, CD8α, TCR | CD4 | 20556631, 34524664 | https://pmc.ncbi.nlm.nih.gov/articles/PMC11115818/; https://pmc.ncbi.nlm.nih.gov/articles/PMC10802914/ |
| cKit+Lineage-Sca1+ Cells (KLS) | KLS | CD117, Sca1 | B220, CD11b, CD19, CD3, Ly6g, TCR, Ter119 | 31282128, 19023891 | https://pmc.ncbi.nlm.nih.gov/articles/PMC6852342/; https://pmc.ncbi.nlm.nih.gov/articles/PMC2640229/ |
| Colony Forming Units - Erythroid (CFU-E) | CFU-E | CD117, CD71 | CD41, Ter119 | 36408979 | https://pmc.ncbi.nlm.nih.gov/articles/PMC12879316/ |
| Common Lymphoid Progenitors (CLP) | CLP | CD127 | B220, CD11b, CD19, CD3, Ly6g, TCR, Ter119 | 31282128, 19023891 | https://pmc.ncbi.nlm.nih.gov/articles/PMC6852342/; https://pmc.ncbi.nlm.nih.gov/articles/PMC2640229/ |
| Common Myeloid Progenitors (CMP) | CMP | CD117, CD34 | B220, CD11b, CD19, CD3, Fcgr, Ly6g, Sca1, TCR, Ter119 | 31282128, 19023891 | https://pmc.ncbi.nlm.nih.gov/articles/PMC6852342/; https://pmc.ncbi.nlm.nih.gov/articles/PMC2640229/ |
| Conventional Dendritic Cells (cDC) | cDC | CD11c, MHC II |  | 31668803 | https://pmc.ncbi.nlm.nih.gov/articles/PMC6838684/ |
| Double Negative T Cells | Double Negative T Cells | CD3, TCR | CD4, CD8α | 34524664 | https://pmc.ncbi.nlm.nih.gov/articles/PMC10802914/ |
| Erythroblasts | Erythroblasts | CD71, Ter119 | CD45 | 21847081 | https://pmc.ncbi.nlm.nih.gov/articles/PMC3211121/ |
| Erythrocytes | Erythrocytes | Ter119 | CD45, CD71 | 21847081 | https://pmc.ncbi.nlm.nih.gov/articles/PMC3211121/ |
| Granulocyte-Macrophage Progenitors (GMP) | GMP | CD117, Fcgr | B220, CD11b, CD19, CD3, Ly6g, Sca1, TCR, Ter119 | 31282128, 19023891 | https://pmc.ncbi.nlm.nih.gov/articles/PMC6852342/; https://pmc.ncbi.nlm.nih.gov/articles/PMC2640229/ |
| Lepr+ Perivascular Cells | Lepr+ Perivascular Cells | Lepr | CD31, CD45, Ter119 | 24953181 | https://pmc.ncbi.nlm.nih.gov/articles/PMC4127103/ |
| Megakaryocyte Progenitors (MkP) | MkP | CD117, CD41 | Ter119 | 30573502, 15779060 | https://haematologica.org/article/view/9052; https://onlinelibrary.wiley.com/doi/10.1002/cyto.a.20133 |
| Megakaryocyte-Erythroid Progenitors (MEP) | MEP | CD117 | B220, CD11b, CD19, CD3, CD34, Fcgr, Ly6g, Sca1, TCR, Ter119 | 31282128 | https://pmc.ncbi.nlm.nih.gov/articles/PMC6852342/ |
| Megakaryocytes | Megakaryocytes | CD41 | CD117, CD45, Ter119 | 30573502, 15779060 | https://haematologica.org/article/view/9052; https://onlinelibrary.wiley.com/doi/10.1002/cyto.a.20133 |
| Monocytes* | Monocytes | CD115, CD11b, F4/80 | Ly6g | 20556631 | https://pmc.ncbi.nlm.nih.gov/articles/PMC11115818/ |
| Neutrophils | Neutrophils | CD11b, Ly6g | F4/80 | 20556631 | https://pmc.ncbi.nlm.nih.gov/articles/PMC11115818/ |
| Other Bone Marrow Endothelial Cells (Other BM ECs) | Other BM ECs | CD105 | CD45, Ter119 | 20556631 | https://pmc.ncbi.nlm.nih.gov/articles/PMC11115818/ |

**Table EV2.**
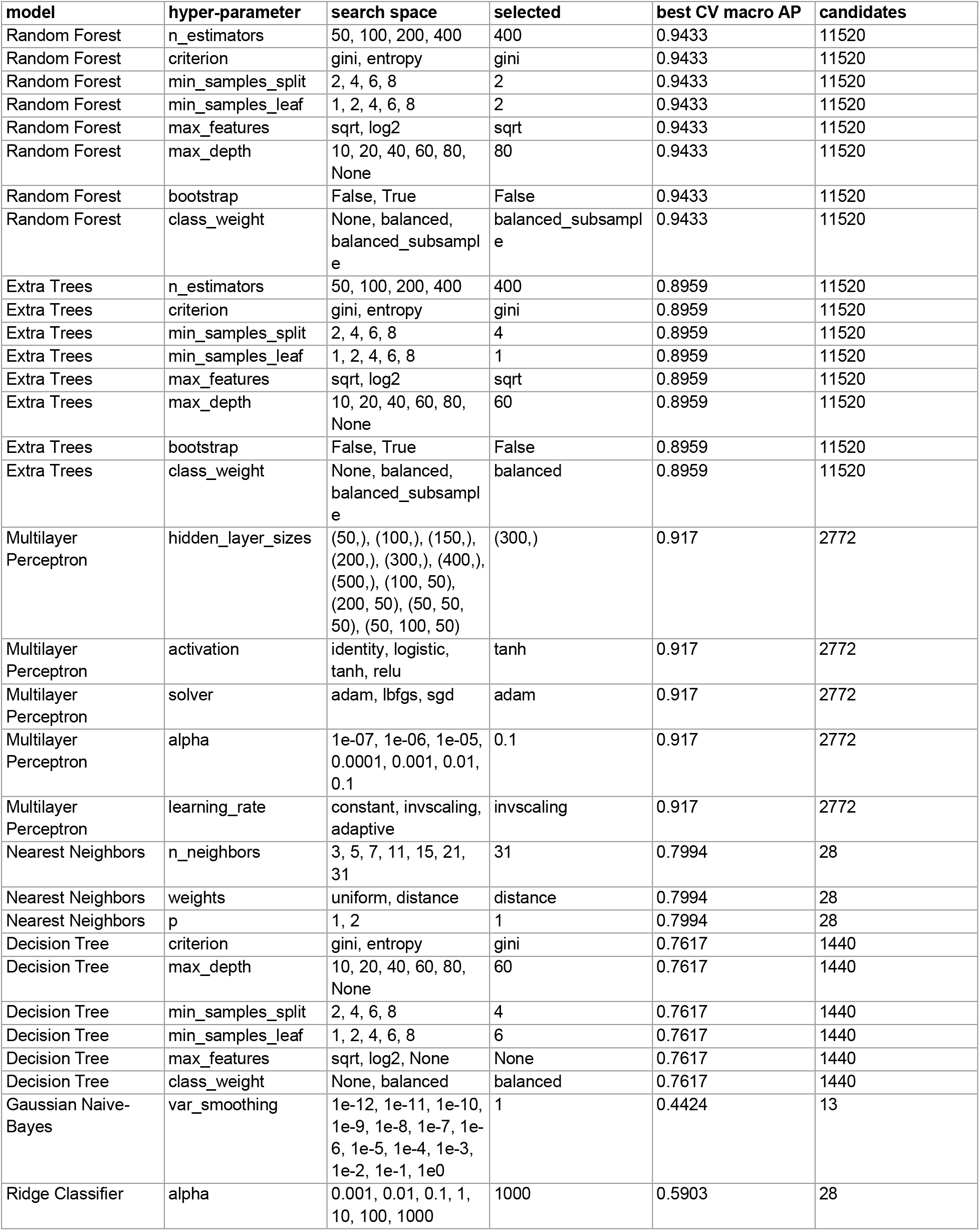
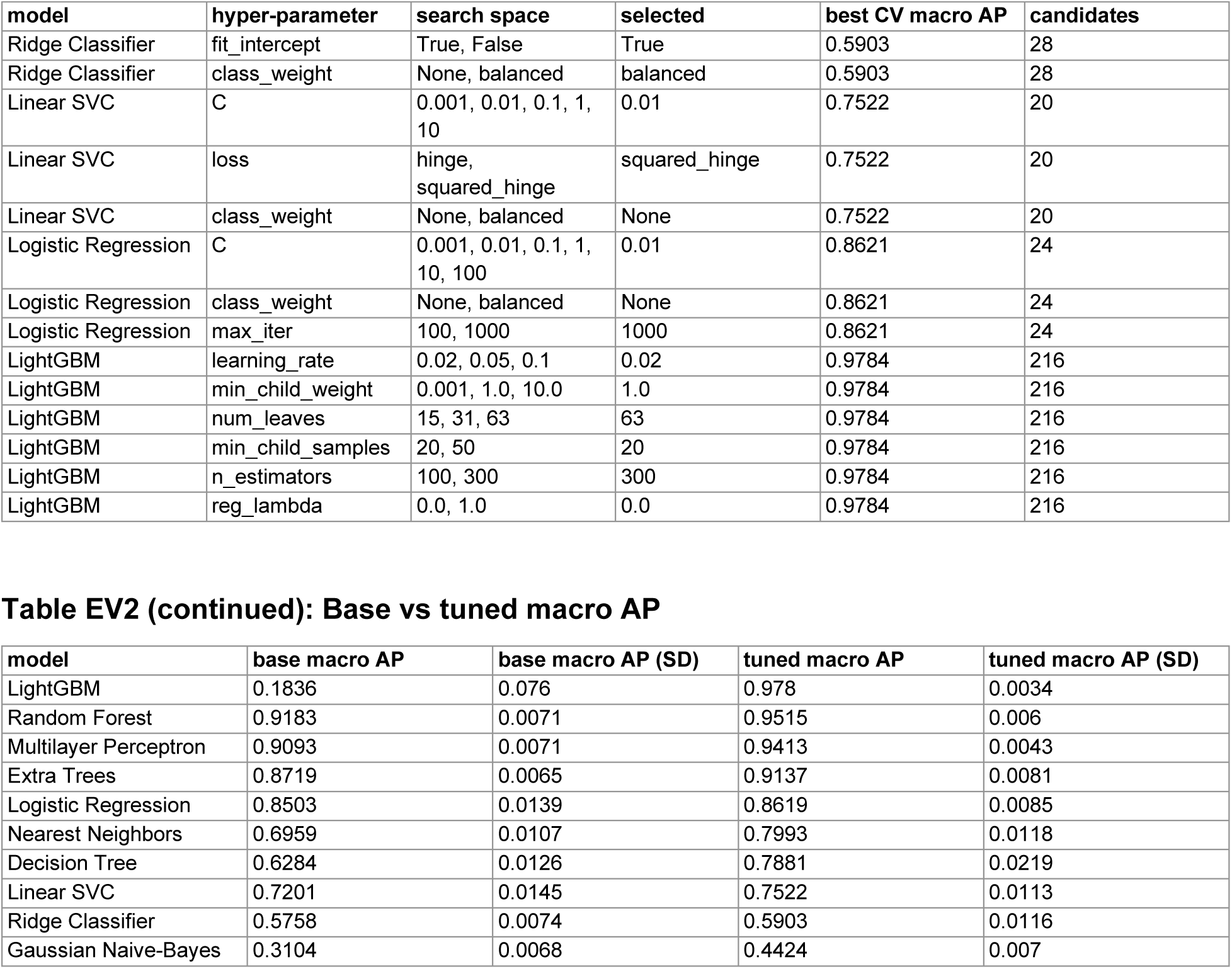
Hyperparameter tuning for model optimization. Hyperparameter tuning for model optimization. Macro AP scores shown are the average across 10-fold cross-validation.

| model | hyper-parameter | search space | selected | best CV macro AP | candidates |
|---|---|---|---|---|---|
| Random Forest | n_estimators | 50, 100, 200, 400 | 400 | 0.9433 | 11520 |
| Random Forest | criterion | gini, entropy | gini | 0.9433 | 11520 |
| Random Forest | min_samples_split | 2, 4, 6, 8 | 2 | 0.9433 | 11520 |
| Random Forest | min_samples_leaf | 1, 2, 4, 6, 8 | 2 | 0.9433 | 11520 |
| Random Forest | max_features | sqrt, log2 | sqrt | 0.9433 | 11520 |
| Random Forest | max_depth | 10, 20, 40, 60, 80, None | 80 | 0.9433 | 11520 |
| Random Forest | bootstrap | False, True | False | 0.9433 | 11520 |
| Random Forest | class_weight | None, balanced, balanced_subsample | balanced_subsample | 0.9433 | 11520 |
| Extra Trees | n_estimators | 50, 100, 200, 400 | 400 | 0.8959 | 11520 |
| Extra Trees | criterion | gini, entropy | gini | 0.8959 | 11520 |
| Extra Trees | min_samples_split | 2, 4, 6, 8 | 4 | 0.8959 | 11520 |
| Extra Trees | min_samples_leaf | 1, 2, 4, 6, 8 | 1 | 0.8959 | 11520 |
| Extra Trees | max_features | sqrt, log2 | sqrt | 0.8959 | 11520 |
| Extra Trees | max_depth | 10, 20, 40, 60, 80, None | 60 | 0.8959 | 11520 |
| Extra Trees | bootstrap | False, True | False | 0.8959 | 11520 |
| Extra Trees | class_weight | None, balanced, balanced_subsample | balanced | 0.8959 | 11520 |
| Multilayer Perceptron | hidden_layer_sizes | (50,), (100,), (150,), (200,), (300,), (400,), (500,), (100, 50), (200, 50), (50, 50, 50), (50, 100, 50) | (300,) | 0.917 | 2772 |
| Multilayer Perceptron | activation | identity, logistic, tanh, relu | tanh | 0.917 | 2772 |
| Multilayer Perceptron | solver | adam, lbfgs, sgd | adam | 0.917 | 2772 |
| Multilayer Perceptron | alpha | 1e-07, 1e-06, 1e-05, 0.0001, 0.001, 0.01, 0.1 | 0.1 | 0.917 | 2772 |
| Multilayer Perceptron | learning_rate | constant, invscaling, adaptive | invscaling | 0.917 | 2772 |
| Nearest Neighbors | n_neighbors | 3, 5, 7, 11, 15, 21, 31 | 31 | 0.7994 | 28 |
| Nearest Neighbors | weights | uniform, distance | distance | 0.7994 | 28 |
| Nearest Neighbors | p | 1, 2 | 1 | 0.7994 | 28 |
| Decision Tree | criterion | gini, entropy | gini | 0.7617 | 1440 |
| Decision Tree | max_depth | 10, 20, 40, 60, 80, None | 60 | 0.7617 | 1440 |
| Decision Tree | min_samples_split | 2, 4, 6, 8 | 4 | 0.7617 | 1440 |
| Decision Tree | min_samples_leaf | 1, 2, 4, 6, 8 | 6 | 0.7617 | 1440 |
| Decision Tree | max_features | sqrt, log2, None | None | 0.7617 | 1440 |
| Decision Tree | class_weight | None, balanced | balanced | 0.7617 | 1440 |
| Gaussian Naive-Bayes | var_smoothing | 1e-12, 1e-11, 1e-10, 1e-9, 1e-8, 1e-7, 1e-6, 1e-5, 1e-4, 1e-3, 1e-2, 1e-1, 1e0 | 1 | 0.4424 | 13 |
| Ridge Classifier | alpha | 0.001, 0.01, 0.1, 1, 10, 100, 1000 | 1000 | 0.5903 | 28 |
| **model** | **hyper-parameter** | **search space** | **selected** | **best CV macro AP** | **candidates** |
| Ridge Classifier | fit_intercept | True, False | True | 0.5903 | 28 |
| Ridge Classifier | class_weight | None, balanced | balanced | 0.5903 | 28 |
| Linear SVC | C | 0.001, 0.01, 0.1, 1, 10 | 0.01 | 0.7522 | 20 |
| Linear SVC | loss | hinge, squared_hinge | squared_hinge | 0.7522 | 20 |
| Linear SVC | class_weight | None, balanced | None | 0.7522 | 20 |
| Logistic Regression | C | 0.001, 0.01, 0.1, 1, 10, 100 | 0.01 | 0.8621 | 24 |
| Logistic Regression | class_weight | None, balanced | None | 0.8621 | 24 |
| Logistic Regression | max_iter | 100, 1000 | 1000 | 0.8621 | 24 |
| LightGBM | learning_rate | 0.02, 0.05, 0.1 | 0.02 | 0.9784 | 216 |
| LightGBM | min_child_weight | 0.001, 1.0, 10.0 | 1.0 | 0.9784 | 216 |
| LightGBM | num_leaves | 15, 31, 63 | 63 | 0.9784 | 216 |
| LightGBM | min_child_samples | 20, 50 | 20 | 0.9784 | 216 |
| LightGBM | n_estimators | 100, 300 | 300 | 0.9784 | 216 |
| LightGBM | reg_lambda | 0.0, 1.0 | 0.0 | 0.9784 | 216 |
| **model** | **base macro AP** | **base macro AP (SD)** | **tuned macro AP** | **tuned macro AP (SD)** |  |
| LightGBM | 0.1836 | 0.076 | 0.978 | 0.0034 |  |
| Random Forest | 0.9183 | 0.0071 | 0.9515 | 0.006 |  |
| Multilayer Perceptron | 0.9093 | 0.0071 | 0.9413 | 0.0043 |  |
| Extra Trees | 0.8719 | 0.0065 | 0.9137 | 0.0081 |  |
| Logistic Regression | 0.8503 | 0.0139 | 0.8619 | 0.0085 |  |
| Nearest Neighbors | 0.6959 | 0.0107 | 0.7993 | 0.0118 |  |
| Decision Tree | 0.6284 | 0.0126 | 0.7881 | 0.0219 |  |
| Linear SVC | 0.7201 | 0.0145 | 0.7522 | 0.0113 |  |
| Ridge Classifier | 0.5758 | 0.0074 | 0.5903 | 0.0116 |  |
| Gaussian Naive-Bayes | 0.3104 | 0.0068 | 0.4424 | 0.007 |  |

**Table EV3.** Additional performance metrics across models and datasets. Additional performance metrics across models and datasets. Results shown are from the same benchmarking as Fig 4. Coverage is the fraction of held-out cells that were assigned a cell type.

| **dataset** | **method** | **held-out cells** | **cell types** | **coverage** | **precision (macro)** | **recall (macro)** | **F1 (macro)** | **accuracy (all cells)** | **accuracy (assigned cells)** |
|---|---|---|---|---|---|---|---|---|---|
| Bone marrow | AnnoSpat | 20154 | 21 | 0.932 | 0.304 | 0.43 | 0.28 | 0.309 | 0.332 |
| Bone marrow | Astir | 20154 | 21 | 0.36 | 0.352 | 0.264 | 0.263 | 0.238 | 0.663 |
| Bone marrow | CELESTA | 20154 | 21 | 0.829 | 0.707 | 0.633 | 0.634 | 0.673 | 0.813 |
| Bone marrow | MAPS | 20154 | 21 | 1 | 0.759 | 0.854 | 0.799 | 0.899 | 0.899 |
| Bone marrow | QuantCell (LightGBM) | 20154 | 21 | 0.994 | 0.967 | 0.871 | 0.901 | 0.967 | 0.973 |
| Colon | AnnoSpat | 48901 | 27 | 0.912 | 0.205 | 0.182 | 0.155 | 0.303 | 0.332 |
| Colon | Astir | 48901 | 27 | 0.226 | 0.266 | 0.151 | 0.138 | 0.159 | 0.702 |
| Colon | CELESTA | 48901 | 27 | 0.689 | 0.25 | 0.215 | 0.195 | 0.376 | 0.546 |
| Colon | MAPS | 48901 | 27 | 1 | 0.706 | 0.869 | 0.765 | 0.855 | 0.855 |
| Colon | QuantCell (random forest) | 48901 | 27 | 0.699 | 0.925 | 0.466 | 0.583 | 0.665 | 0.95 |
| Lymph node (cHL1) | AnnoSpat | 307687 | 13 | 0.7 | 0.368 | 0.366 | 0.311 | 0.302 | 0.432 |
| Lymph node (cHL1) | Astir | 307687 | 13 | 0.675 | 0.487 | 0.26 | 0.297 | 0.188 | 0.279 |
| Lymph node (cHL1) | CELESTA | 307687 | 13 | 0.001 | 0.33 | 0.005 | 0.009 | 0.001 | 0.56 |
| Lymph node (cHL1) | MAPS | 307687 | 13 | 1 | 0.86 | 0.943 | 0.896 | 0.916 | 0.916 |
| Lymph node (cHL1) | QuantCell (LightGBM) | 307687 | 13 | 0.954 | 0.954 | 0.897 | 0.921 | 0.912 | 0.955 |
| Lymph node (cHL CODEX) | AnnoSpat | 27725 | 16 | 0.948 | 0.514 | 0.499 | 0.486 | 0.546 | 0.576 |
| Lymph node (cHL CODEX) | Astir | 27725 | 16 | 0.623 | 0.462 | 0.384 | 0.399 | 0.397 | 0.637 |
| Lymph node (cHL CODEX) | CELESTA | 27725 | 16 | 0.301 | 0.548 | 0.251 | 0.269 | 0.169 | 0.56 |
| Lymph node (cHL CODEX) | MAPS | 27725 | 16 | 1 | 0.795 | 0.883 | 0.83 | 0.847 | 0.847 |
| Lymph node (cHL CODEX) | QuantCell (LightGBM) | 27725 | 16 | 0.863 | 0.954 | 0.797 | 0.864 | 0.82 | 0.951 |
| Lymph node (cHL2) | AnnoSpat | 43027 | 11 | 0.762 | 0.528 | 0.369 | 0.397 | 0.358 | 0.47 |
| Lymph node (cHL2) | Astir | 43027 | 11 | 0.309 | 0.394 | 0.116 | 0.167 | 0.174 | 0.562 |
| Lymph node (cHL2) | CELESTA | 43027 | 11 | 0.001 | 0.181 | 0.001 | 0.003 | 0.001 | 0.667 |
| Lymph node (cHL2) | MAPS | 43027 | 11 | 1 | 0.85 | 0.905 | 0.874 | 0.876 | 0.876 |
| Lymph node (cHL2) | QuantCell (LightGBM) | 43027 | 11 | 0.879 | 0.95 | 0.833 | 0.885 | 0.835 | 0.95 |

**Table EV4.** Performance metrics separated by cell frequency across models and datasets. Performance metrics separated by cell frequency across models and datasets. Results shown are from the same benchmarking as Fig 4. Frequencies are based on the held-out ground truth.

| **dataset** | **method** | **cell types below 0.1 frequency** | **precision (<0.1)** | **recall (<0.1)** | **F1 (<0.1)** | **cell types at or above 0.1 frequency** | **precision (≥0.1)** | **recall (≥0.1)** | **F1 (≥0.1)** |
|---|---|---|---|---|---|---|---|---|---|
| Bone marrow | AnnoSpat | 17 | 0.228 | 0.472 | 0.263 | 4 | 0.626 | 0.251 | 0.35 |
| Bone marrow | Astir | 17 | 0.296 | 0.272 | 0.264 | 4 | 0.592 | 0.228 | 0.259 |
| Bone marrow | CELESTA | 17 | 0.681 | 0.636 | 0.631 | 4 | 0.821 | 0.621 | 0.649 |
| Bone marrow | MAPS | 17 | 0.719 | 0.846 | 0.774 | 4 | 0.925 | 0.888 | 0.905 |
| Bone marrow | QuantCell (LightGBM) | 17 | 0.966 | 0.847 | 0.885 | 4 | 0.97 | 0.971 | 0.971 |
| Colon | AnnoSpat | 24 | 0.132 | 0.155 | 0.114 | 3 | 0.792 | 0.391 | 0.485 |
| Colon | Astir | 24 | 0.181 | 0.147 | 0.119 | 3 | 0.951 | 0.183 | 0.297 |
| Colon | CELESTA | 24 | 0.19 | 0.181 | 0.153 | 3 | 0.724 | 0.487 | 0.535 |
| Colon | MAPS | 24 | 0.676 | 0.87 | 0.748 | 3 | 0.941 | 0.858 | 0.898 |
| Colon | QuantCell (random forest) | 24 | 0.921 | 0.424 | 0.547 | 3 | 0.95 | 0.798 | 0.867 |
| Lymph node (cHL1) | AnnoSpat | 9 | 0.254 | 0.412 | 0.288 | 4 | 0.627 | 0.263 | 0.362 |
| Lymph node (cHL1) | Astir | 9 | 0.454 | 0.301 | 0.321 | 4 | 0.561 | 0.167 | 0.245 |
| Lymph node (cHL1) | CELESTA | 9 | 0.232 | 0.007 | 0.013 | 4 | 0.551 | 0 | 0 |
| Lymph node (cHL1) | MAPS | 9 | 0.821 | 0.96 | 0.883 | 4 | 0.946 | 0.906 | 0.925 |
| Lymph node (cHL1) | QuantCell (LightGBM) | 9 | 0.953 | 0.881 | 0.911 | 4 | 0.955 | 0.933 | 0.944 |
| Lymph node (cHL CODEX) | AnnoSpat | 13 | 0.456 | 0.46 | 0.436 | 3 | 0.764 | 0.666 | 0.7 |
| Lymph node (cHL CODEX) | Astir | 13 | 0.398 | 0.368 | 0.364 | 3 | 0.737 | 0.453 | 0.551 |
| Lymph node (cHL CODEX) | CELESTA | 13 | 0.606 | 0.269 | 0.29 | 3 | 0.296 | 0.173 | 0.178 |
| Lymph node (cHL CODEX) | MAPS | 13 | 0.77 | 0.893 | 0.822 | 3 | 0.9 | 0.837 | 0.867 |
| Lymph node (cHL CODEX) | QuantCell (LightGBM) | 13 | 0.954 | 0.786 | 0.857 | 3 | 0.951 | 0.849 | 0.896 |
| Lymph node (cHL2) | AnnoSpat | 7 | 0.461 | 0.379 | 0.395 | 4 | 0.644 | 0.351 | 0.402 |
| Lymph node (cHL2) | Astir | 7 | 0.241 | 0.052 | 0.076 | 4 | 0.661 | 0.227 | 0.326 |
| Lymph node (cHL2) | CELESTA | 7 | 0.284 | 0.002 | 0.005 | 4 | 0 | 0 | 0 |
| Lymph node (cHL2) | MAPS | 7 | 0.816 | 0.923 | 0.865 | 4 | 0.909 | 0.873 | 0.89 |
| Lymph node (cHL2) | QuantCell (LightGBM) | 7 | 0.95 | 0.812 | 0.871 | 4 | 0.95 | 0.871 | 0.908 |

**Appendix Figure S1.**
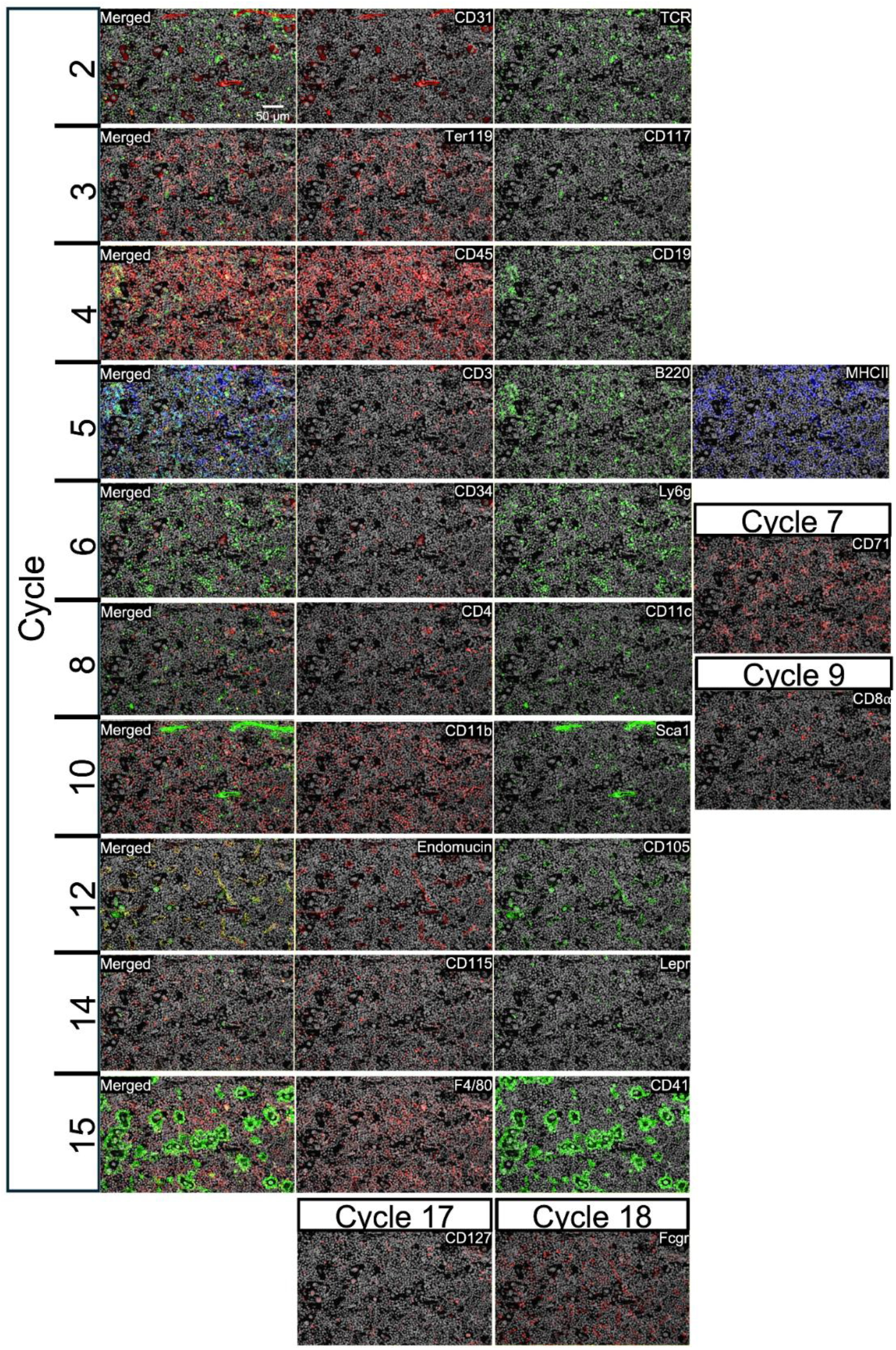
Protein marker staining across PhenoCycler imaging cycles. The first column shows the merged images of all markers imaged within each cycle. Subsequent columns display the individual markers from that cycle. Cycles not shown were either intentionally left blank or contained antibodies that were not included in the final analysis. Cycles 7, 9, 17, and 18 contained only a single marker. DAPI (shown in gray) is included in all images. All images were acquired at 20X magnification. Scale bar, 50 µm.

**Appendix Figure S2.**
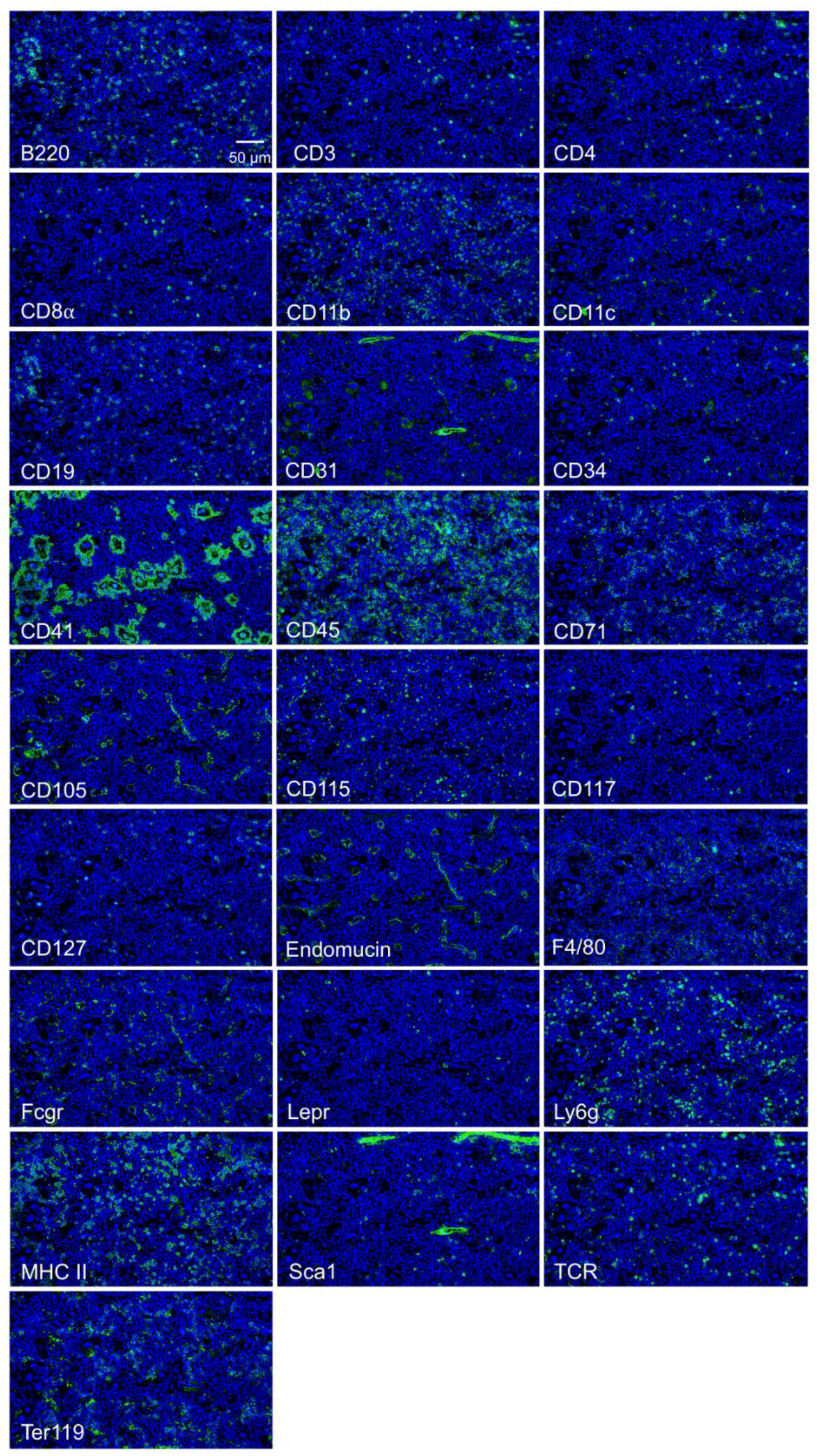
Single channel protein marker staining used in the final analysis. Green indicates the protein marker signal, and blue represents DAPI. All images were acquired at 20X magnification. Scale bar, 50 µm.

**Appendix Figure S3.**
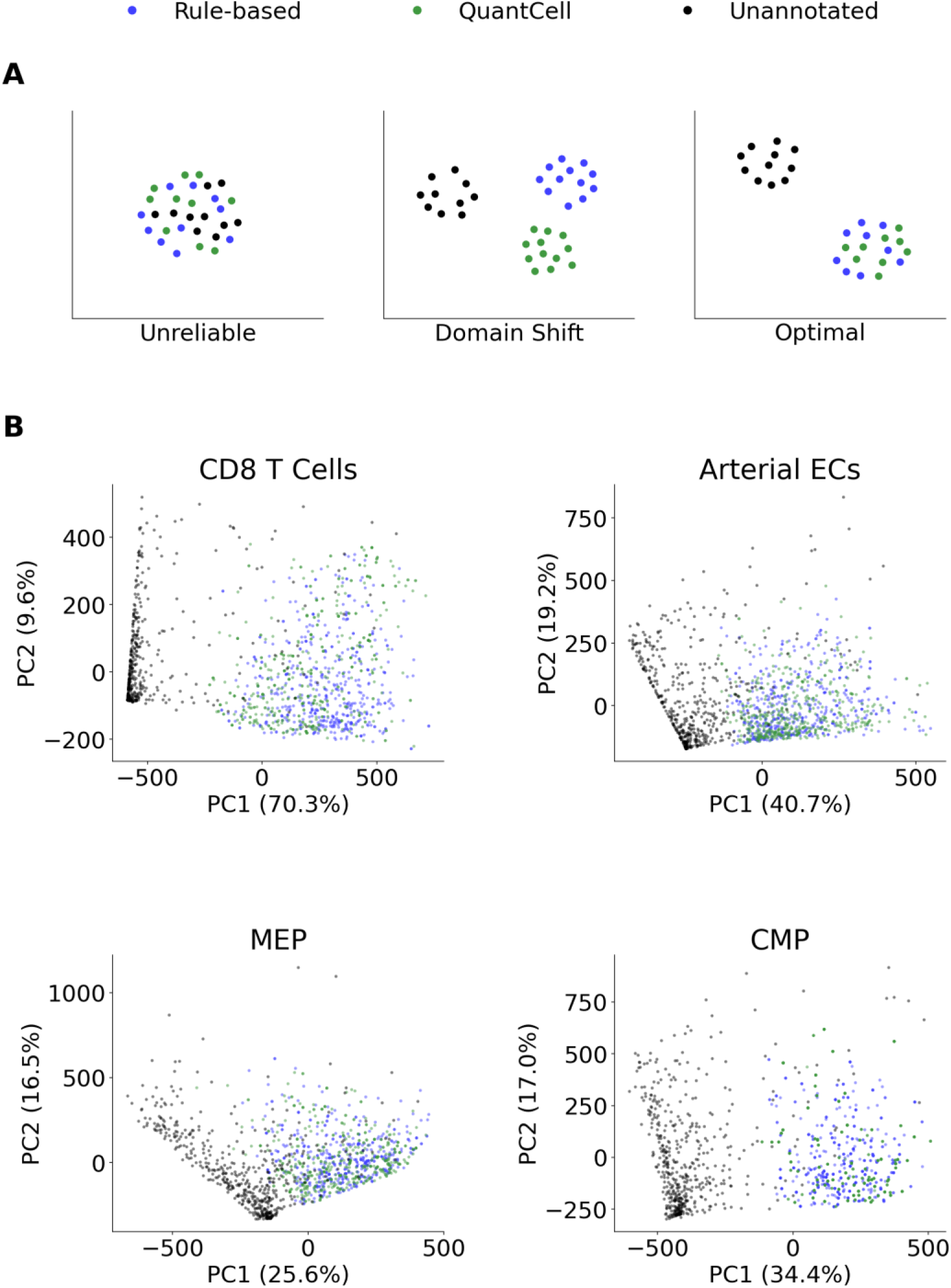
Distribution of binary marker rule-based annotated and QuantCell-annotated cells in principal component space. **(A)** Diagram illustrating the interpretation of representative results. (B) Each panel shows 500 randomly selected cells from the rule-based annotation, QuantCell annotation, and the set of cells not assigned to the corresponding cell type by either method (unannotated). The first two principal components are plotted after performing PCA on quantitative marker data using only the markers used for the rule-based annotation of that cell type.

**Appendix Figure S4.**
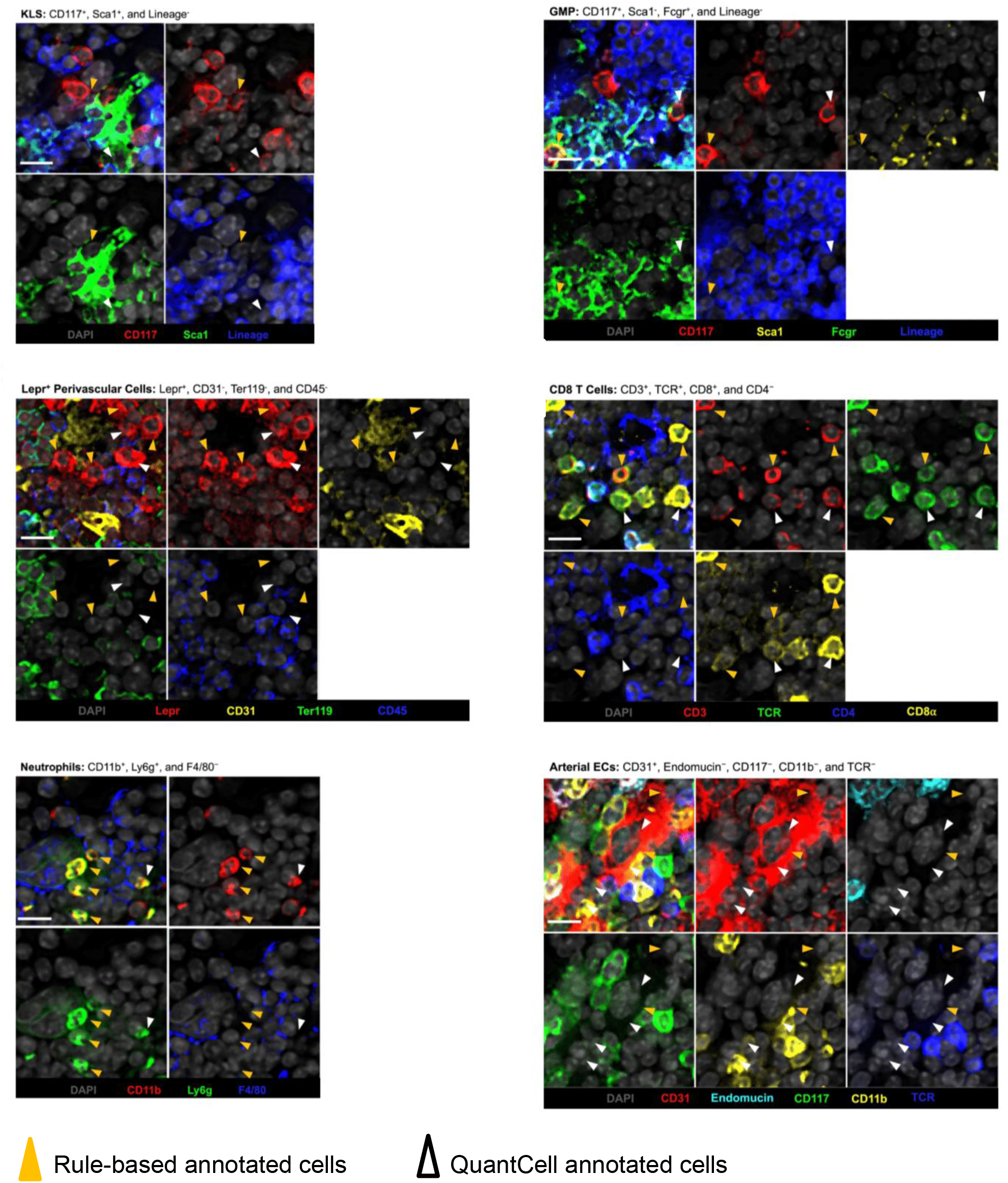
Representative images of cells identified by rule-based annotation and by QuantCell. Merged images (top left panel) and individual marker stains are shown for the representative cell types. Scale bar is 10 µm. Lineage is a composite image of Ter119, Ly6g, CD11b, TCR, B220, CD19, and CD3. All images were acquired at 20X magnification.

**Appendix Figure S5.**
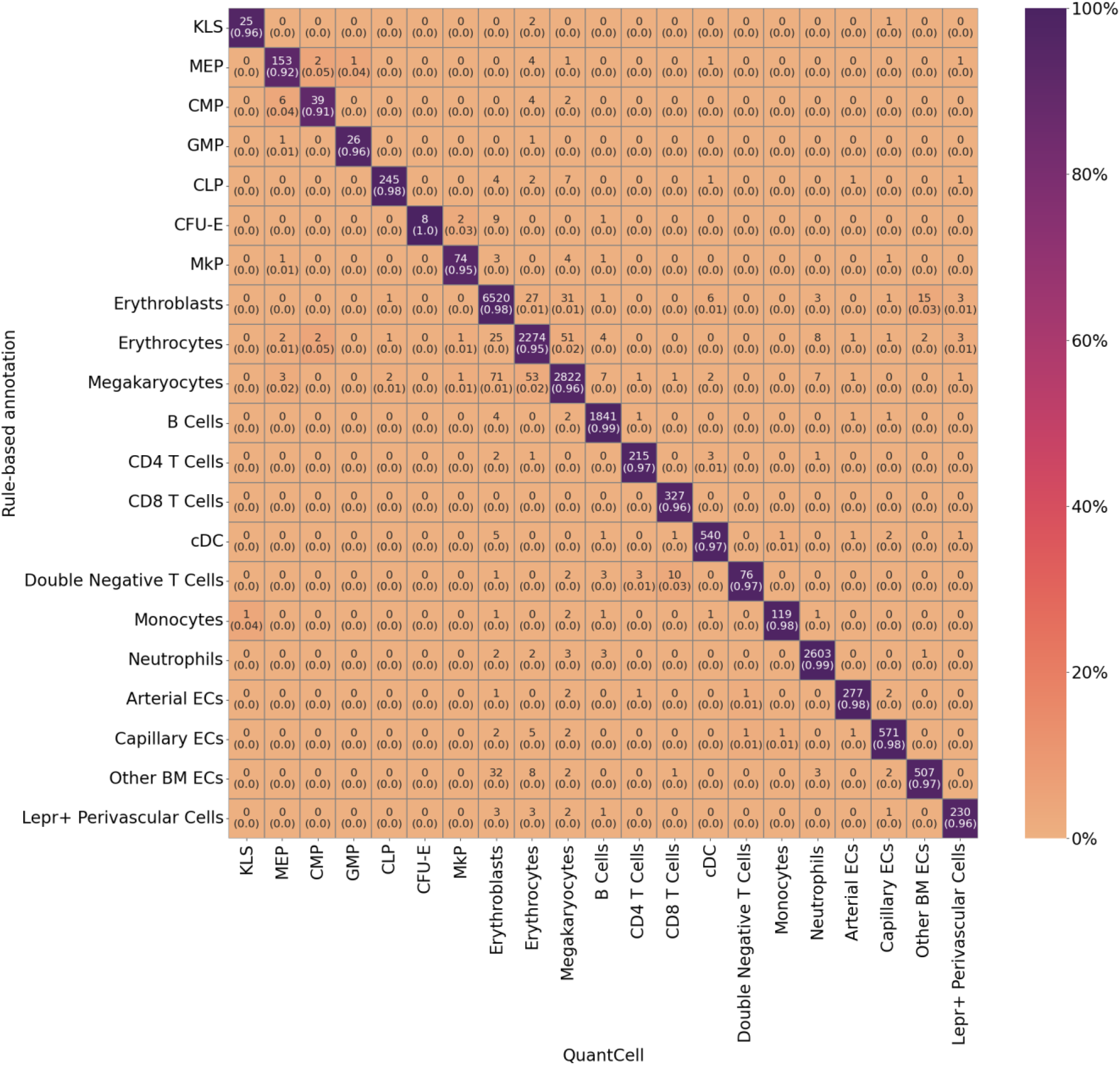
Confusion matrix for QuantCell performance. The matrix is column-normalized and compares rule-based cell annotation with QuantCell annotation at a 5% FDR threshold. Diagonal values correspond to precision. Each cell displays the raw count and the column-normalized fraction. n = 20,038 annotated test cells.

**Appendix Figure S6.**
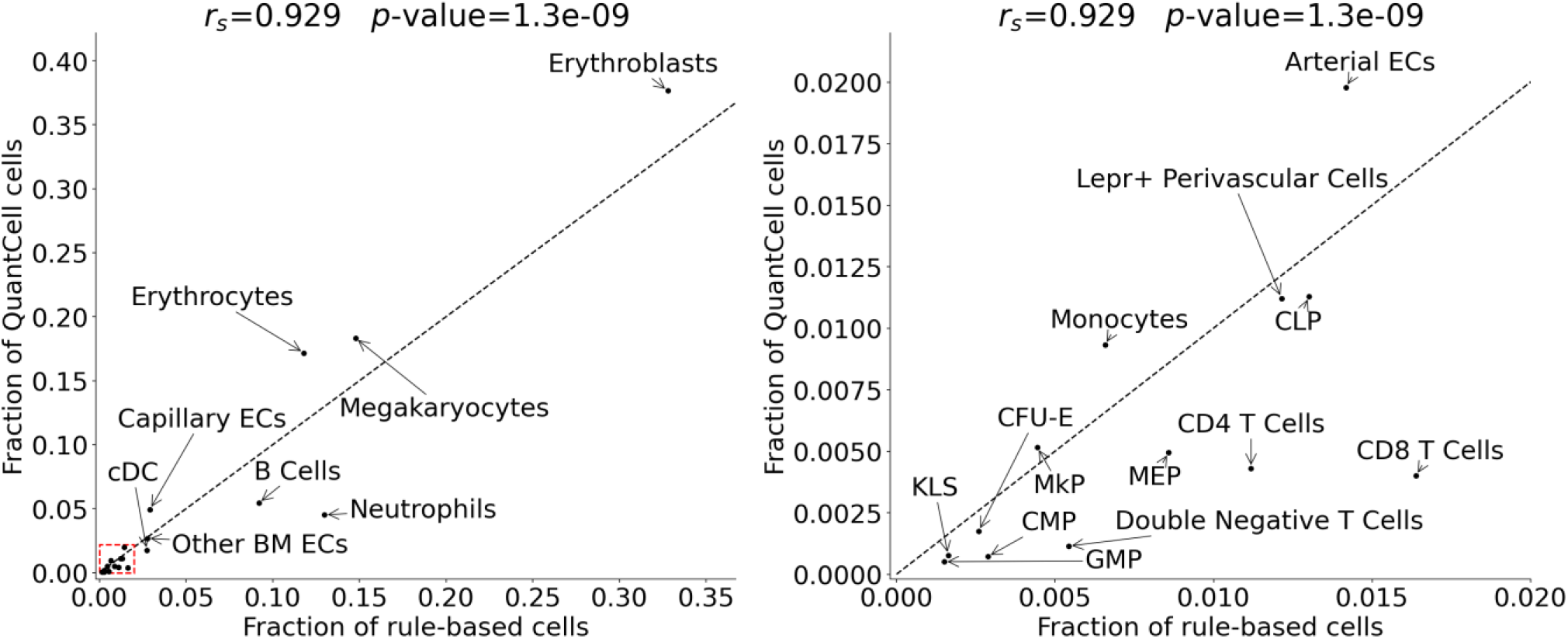
Comparison of cell type frequencies obtained from rule-based annotation and QuantCell annotation. Data are presented as fractions of cells annotated by that method. Each dot represents one cell type (n = 21). The graph on the right shows a magnified view of the red dashed region in the left panel. The dashed line represents equivalence.

**Appendix Figure S7.**
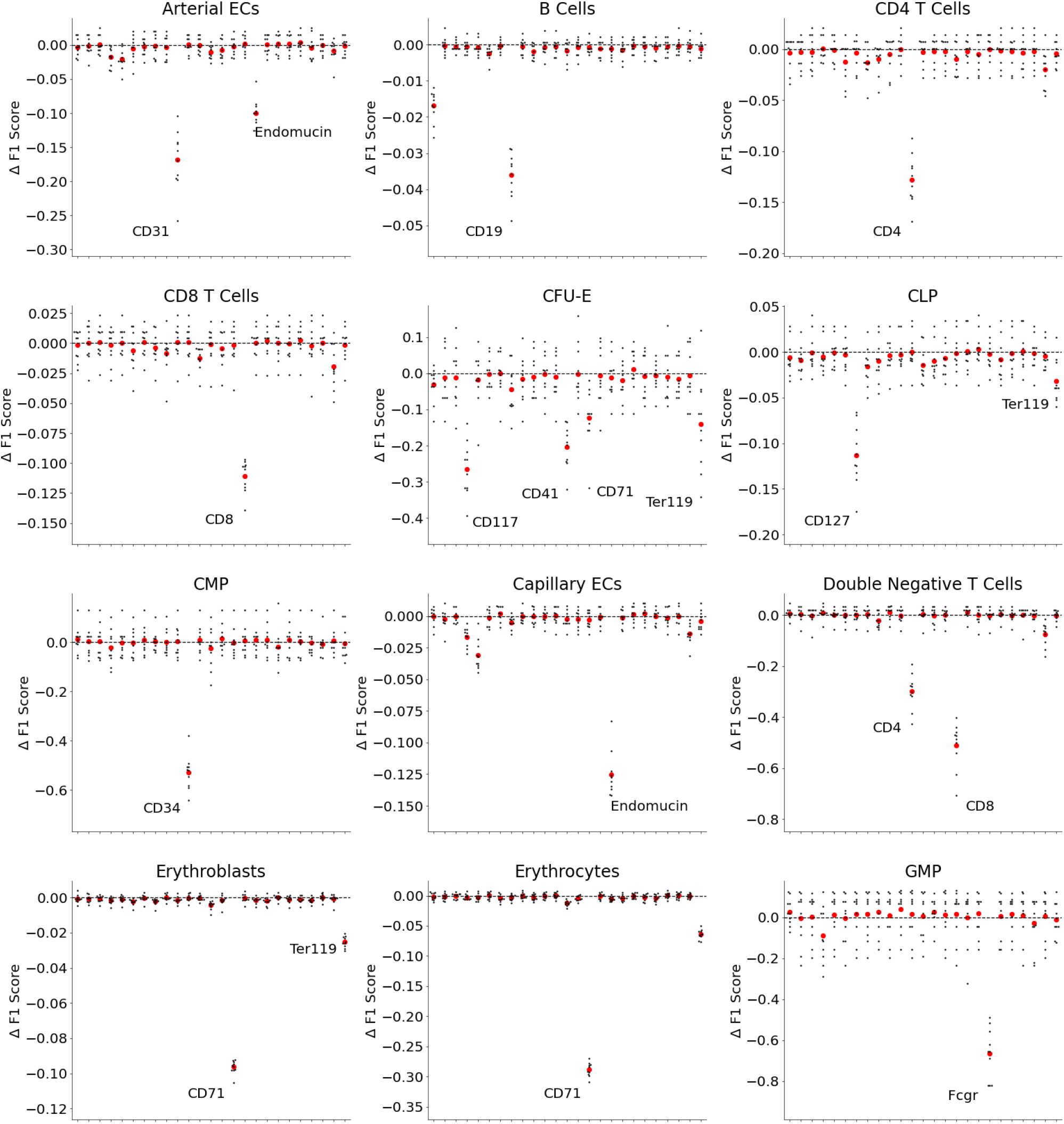
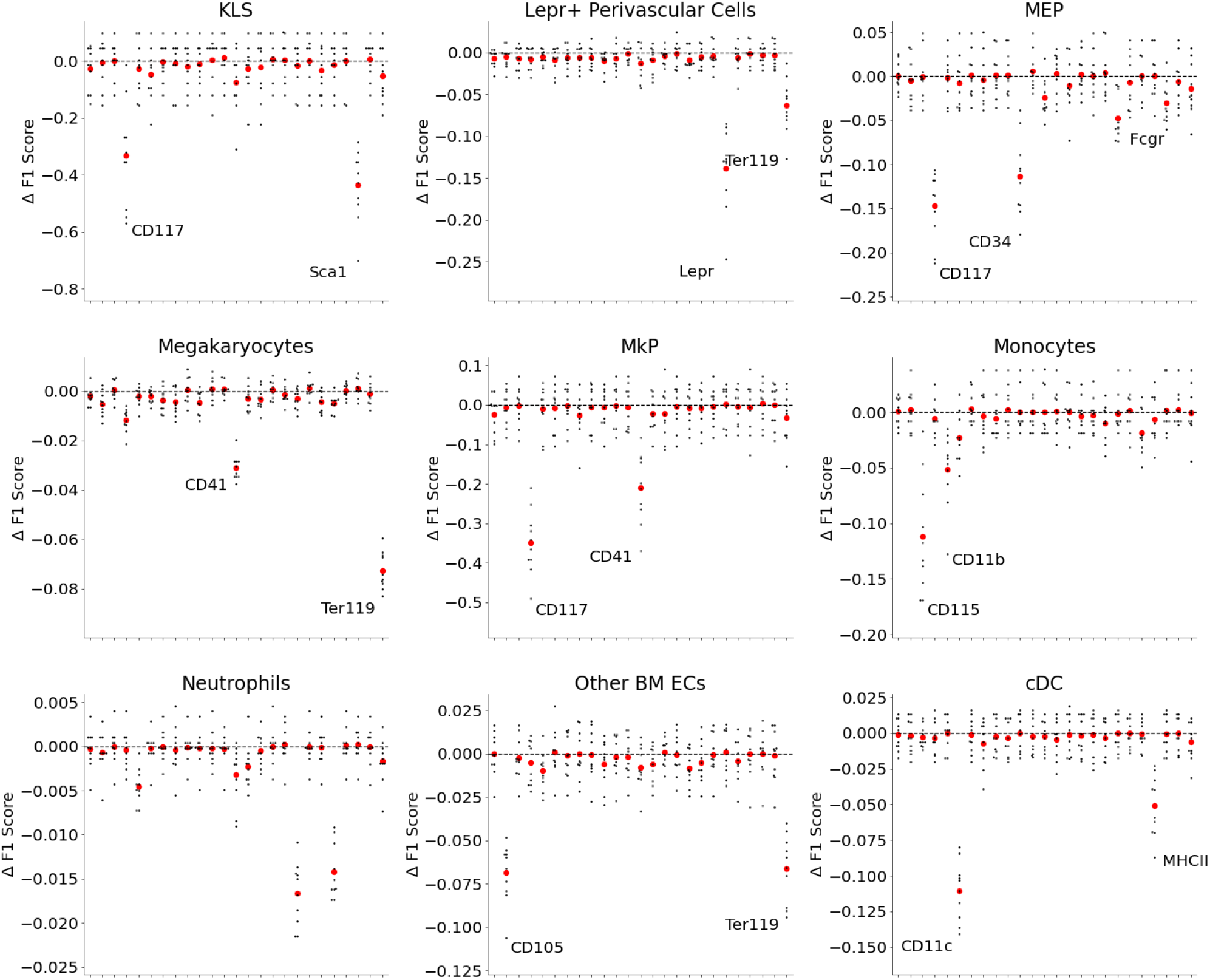
Impact of individual protein markers on prediction performance for additional cell types. Shown are changes in F1 score resulting from the exclusion of individual protein markers from the training and testing datasets. Each dot represents the change in the F1 score relative to the baseline (no exclusion), calculated using 10-fold cross-validation. Red dots indicate the mean F1 score change across all 10 folds. Prominent markers are labeled.

## References

Abbaszadegan MR, Bagheri V, Razavi MS, Momtazi AA, Sahebkar A, Gholamin M (2017) Isolation, identification, and characterization of cancer stem cells: A review. J Cell Physiol 232: 2008–2018

Altshuler A, Wickström SA, Shalom-Feuerstein R (2023) Spotlighting adult stem cells: advances, pitfalls, and challenges. Trends Cell Biol 33: 477–494

Amitay Y, Bussi Y, Feinstein B, Bagon S, Milo I, Keren L (2023) CellSighter: a neural network to classify cells in highly multiplexed images. Nat Commun 14: 4302

Baccin C, Al-Sabah J, Velten L, Helbling PM, Grünschläger F, Hernández-Malmierca P, Nombela-Arrieta C, Steinmetz LM, Trumpp A, Haas S (2020) Combined single-cell and spatial transcriptomics reveal the molecular, cellular and spatial bone marrow niche organization. Nat Cell Biol 22: 38–48

Bae S, Na KJ, Koh J, Lee DS, Choi H, Kim YT (2022) CellDART: cell type inference by domain adaptation of single-cell and spatial transcriptomic data. Nucleic Acids Res 50: e57

Bai Y, Zhu B, Rovira-Clave X, Chen H, Markovic M, Chan CN, Su T-H, McIlwain DR, Estes JD, Keren L et al (2021) Adjacent cell marker lateral spillover compensation and reinforcement for multiplexed images. Front Immunol 12: 652631

Bankhead P, Loughrey MB, Fernández JA, Dombrowski Y, McArt DG, Dunne PD, McQuaid S, Gray RT, Murray LJ, Coleman HG et al (2017) QuPath: Open source software for digital pathology image analysis. Sci Rep 7: 16878

Black S, Phillips D, Hickey JW, Kennedy-Darling J, Venkataraaman VG, Samusik N, Goltsev Y, Schürch CM, Nolan GP (2021) CODEX multiplexed tissue imaging with DNA-conjugated antibodies. Nat Protoc 16: 3802–3835

Blise KE, Sivagnanam S, Banik GL, Coussens LM, Goecks J (2022) Single-cell spatial architectures associated with clinical outcome in head and neck squamous cell carcinoma. npj Precis Oncol 6: 10

Boohar W, Wang B, Thomas Z, Nogalska A, Lu R (2026) Data for QuantCell: machine learning-based cell annotation leveraging quantitative imaging profiles. Zenodo doi:10.5281/zenodo.23672666 (https://doi.org/10.5281/zenodo.23672666) [DATASET]

Boulesteix A-L, Janitza S, Kruppa J, König IR (2012) Overview of random forest methodology and practical guidance with emphasis on computational biology and bioinformatics. WIREs Data Min Knowl Discov 2: 493–507

Breiman L (2001) Random forests. Mach Learn 45: 5–32

Bressan D, Battistoni G, Hannon GJ (2023) The dawn of spatial omics. Science 381: eabq4964

Bryder D, Rossi DJ, Weissman IL (2006) Hematopoietic stem cells: the paradigmatic tissue-specific stem cell. Am J Pathol 169: 338–346

Chen A, Liao S, Cheng M, Ma K, Wu L, Lai Y, Qiu X, Yang J, Xu J, Hao S et al (2022) Spatiotemporal transcriptomic atlas of mouse organogenesis using DNA nanoball-patterned arrays. Cell 185: 1777–1792.e21

Chen KH, Boettiger AN, Moffitt JR, Wang S, Zhuang X (2015) Spatially resolved, highly multiplexed RNA profiling in single cells. Science 348: aaa6090

Chen X, Ishwaran H (2012) Random forests for genomic data analysis. Genomics 99: 323–329

Cheng X, Peng T, Chu T, Yang Y, Liu J, Gao Q, Cao C, Wei J (2025) Application of single-cell and spatial omics in deciphering cellular hallmarks of cancer drug response and resistance. J Hematol Oncol 18: 70

Coleman K, Hu J, Schroeder A, Lee EB, Li M (2023) SpaDecon: cell-type deconvolution in spatial transcriptomics with semi-supervised learning. Commun Biol 6: 378

Cossarizza A, Chang H-D, Radbruch A, Acs A, Adam D, Adam-Klages S, Agace WW, Aghaeepour N, Akdis M, Allez M et al (2019) Guidelines for the use of flow cytometry and cell sorting in immunological studies (second edition). Eur J Immunol 49: 1457–1973

Ding L, Morrison SJ (2013) Haematopoietic stem cells and early lymphoid progenitors occupy distinct bone marrow niches. Nature 495: 231–235

Ding L, Saunders TL, Enikolopov G, Morrison SJ (2012) Endothelial and perivascular cells maintain haematopoietic stem cells. Nature 481: 457–462

Dolgalev I, Tikhonova AN (2021) Connecting the dots: resolving the bone marrow niche heterogeneity. Front Cell Dev Biol 9: 622519

Dong K, Zhang S (2022) Deciphering spatial domains from spatially resolved transcriptomics with an adaptive graph attention auto-encoder. Nat Commun 13: 1739

Eng C-HL, Lawson M, Zhu Q, Dries R, Koulena N, Takei Y, Yun J, Cronin C, Karp C, Yuan G-C et al (2019) Transcriptome-scale super-resolved imaging in tissues by RNA seqFISH. Nature 568: 235–239

Ge S, Sun S, Xu H, Cheng Q, Ren Z (2025) Deep learning in single-cell and spatial transcriptomics data analysis: advances and challenges from a data science perspective. Brief Bioinform 26: bbaf136

Geuenich MJ, Hou J, Lee S, Ayub S, Jackson HW, Campbell KR (2021) Automated assignment of cell identity from single-cell multiplexed imaging and proteomic data. Cell Syst 12: 1173–1186.e5

Goltsev Y, Samusik N, Kennedy-Darling J, Bhate S, Hale M, Vazquez G, Black S, Nolan GP (2018) Deep profiling of mouse splenic architecture with CODEX multiplexed imaging. Cell 174: 968–981.e15

Hung M-C, Link W (2011) Protein localization in disease and therapy. J Cell Sci 124: 3381–3392

Ianevski A, Giri AK, Aittokallio T (2022) Fully-automated and ultra-fast cell-type identification using specific marker combinations from single-cell transcriptomic data. Nat Commun 13: 1246

Jamieson K, Talwalkar A (2016) Non-stochastic best arm identification and hyperparameter optimization. In: Proceedings of the 19th International Conference on Artificial Intelligence and Statistics, Gretton A, Robert CC (eds), Proceedings of Machine Learning Research 51, pp 240–248. PMLR

Ke G, Meng Q, Finley T, Wang T, Chen W, Ma W, Ye Q, Liu T-Y (2017) LightGBM: a highly efficient gradient boosting decision tree. In: Advances in Neural Information Processing Systems 30, Guyon I, Von Luxburg U, Bengio S, Wallach H, Fergus R, Vishwanathan S, Garnett R (eds). Curran Associates, Red Hook, NY

Kiselev VY, Andrews TS, Hemberg M (2019) Challenges in unsupervised clustering of single-cell RNA-seq data. Nat Rev Genet 20: 273–282

Kumagai Y (2024) BootCellNet, a resampling-based procedure, promotes unsupervised identification of cell populations via robust inference of gene regulatory networks. PLoS Comput Biol 20: e1012480

Li B, Zhang W, Guo C, Xu H, Li L, Fang M, Hu Y, Zhang X, Yao X, Tang M et al (2022) Benchmarking spatial and single-cell transcriptomics integration methods for transcript distribution prediction and cell type deconvolution. Nat Methods 19: 662–670

Loughrey MB, Bankhead P, Coleman HG, Hagan RS, Craig S, McCorry AMB, Gray RT, McQuaid S, Dunne PD, Hamilton PW et al (2018) Validation of the systematic scoring of immunohistochemically stained tumour tissue microarrays using QuPath digital image analysis. Histopathology 73: 327–338

Lubeck E, Coskun AF, Zhiyentayev T, Ahmad M, Cai L (2014) Single-cell in situ RNA profiling by sequential hybridization. Nat Methods 11: 360–361

Lundberg E, Borner GHH (2019) Spatial proteomics: a powerful discovery tool for cell biology. Nat Rev Mol Cell Biol 20: 285–302

Marghi Y, Gala R, Baftizadeh F, Sümbül U (2024) Joint inference of discrete cell types and continuous type-specific variability in single-cell datasets with MMIDAS. Nat Comput Sci 4: 706–722

Marx V (2021) Method of the Year: spatially resolved transcriptomics. Nat Methods 18: 9–14

Merritt CR, Ong GT, Church SE, Barker K, Danaher P, Geiss G, Hoang M, Jung J, Liang Y, McKay-Fleisch J et al (2020) Multiplex digital spatial profiling of proteins and RNA in fixed tissue. Nat Biotechnol 38: 586–599

Mongia A, Zohora FT, Burget NG, Zhou Y, Saunders DC, Wang YJ, Brissova M, Powers AC, Kaestner KH, Vahedi G et al (2024) AnnoSpat annotates cell types and quantifies cellular arrangements from spatial proteomics. Nat Commun 15: 3744

National Research Council (2011) Guide for the Care and Use of Laboratory Animals, 8th edn. National Academies Press, Washington, DC

Nguyen A, Khoo WH, Moran I, Croucher PI, Phan TG (2018) Single cell RNA sequencing of rare immune cell populations. Front Immunol 9: 1553

Nombela-Arrieta C, Manz MG (2017) Quantification and three-dimensional microanatomical organization of the bone marrow. Blood Adv 1: 407–416

Pachitariu M, Rariden M, Stringer C (2025) Cellpose-SAM: superhuman generalization for cellular segmentation. bioRxiv doi:10.1101/2025.04.28.651001 [PREPRINT]

Pan SJ, Yang Q (2010) A survey on transfer learning. IEEE Trans Knowl Data Eng 22: 1345–1359

Patel SB, Franceski AM, Crown BL, Welner RS (2024) Leading edge techniques in the quest for characterizing rare hematopoietic stem cells. Curr Stem Cell Rep 10: 108–125

Pedregosa F, Varoquaux G, Gramfort A, Michel V, Thirion B, Grisel O, Blondel M, Prettenhofer P, Weiss R, Dubourg V et al (2011) Scikit-learn: machine learning in Python. J Mach Learn Res 12: 2825–2830

Purton LE (2022) Adult murine hematopoietic stem cells and progenitors: an update on their identities, functions, and assays. Exp Hematol 116: 1–14

Rodriques SG, Stickels RR, Goeva A, Martin CA, Murray E, Vanderburg CR, Welch J, Chen LM, Chen F, Macosko EZ (2019) Slide-seq: A scalable technology for measuring genome-wide expression at high spatial resolution. Science 363: 1463–1467

Schmidt U, Weigert M, Broaddus C, Myers G (2018) Cell detection with star-convex polygons. In: Medical Image Computing and Computer Assisted Intervention – MICCAI 2018, Frangi AF, Schnabel JA, Davatzikos C, Alberola-López C, Fichtinger G (eds), pp 265–273. Springer International Publishing, Cham

Schürch C (2020) Coordinated cellular neighborhoods orchestrate antitumoral immunity at the colorectal cancer invasive front (single-cell data table). Mendeley Data doi:10.17632/mpjzbtfgfr.1 (https://doi.org/10.17632/mpjzbtfgfr.1) [DATASET]

Schürch CM, Bhate SS, Barlow GL, Phillips DJ, Noti L, Zlobec I, Chu P, Black S, Demeter J, McIlwain DR et al (2020) Coordinated cellular neighborhoods orchestrate antitumoral immunity at the colorectal cancer invasive front. Cell 182: 1341–1359.e19

Seydel C (2025) Beyond cell atlases: spatial biology reveals mechanisms behind disease. Nat Biotechnol 43: 841–844

Shaban M, Bai Y, Qiu H, Mao S, Yeung J, Yeo YY, Shanmugam V, Chen H, Zhu B, Weirather JL et al (2023) Data for MAPS: pathologist-level cell type annotation from tissue images through machine learning. Zenodo doi:10.5281/zenodo.10067010 (https://doi.org/10.5281/zenodo.10067010) [DATASET]

Shaban M, Bai Y, Qiu H, Mao S, Yeung J, Yeo YY, Shanmugam V, Chen H, Zhu B, Weirather JL et al (2024) MAPS: pathologist-level cell type annotation from tissue images through machine learning. Nat Commun 15: 28

Spangrude GJ, Heimfeld S, Weissman IL (1988) Purification and characterization of mouse hematopoietic stem cells. Science 241: 58–62

Stadler C, Rexhepaj E, Singan VR, Murphy RF, Pepperkok R, Uhlén M, Simpson JC, Lundberg E (2013) Immunofluorescence and fluorescent-protein tagging show high correlation for protein localization in mammalian cells. Nat Methods 10: 315–323

Ståhl PL, Salmén F, Vickovic S, Lundmark A, Navarro JF, Magnusson J, Giacomello S, Asp M, Westholm JO, Huss M et al (2016) Visualization and analysis of gene expression in tissue sections by spatial transcriptomics. Science 353: 78–82

Virtanen P, Gommers R, Oliphant TE, Haberland M, Reddy T, Cournapeau D, Burovski E, Peterson P, Weckesser W, Bright J et al (2020) SciPy 1.0: fundamental algorithms for scientific computing in Python. Nat Methods 17: 261–272

Weber LM, Robinson MD (2016) Comparison of clustering methods for high-dimensional single-cell flow and mass cytometry data. Cytometry A 89: 1084–1096

Wu M, Tao H, Xu T, Zheng X, Wen C, Wang G, Peng Y, Dai Y (2024) Spatial proteomics: unveiling the multidimensional landscape of protein localization in human diseases. Proteome Sci 22: 7

Zhang W, Li I, Reticker-Flynn NE, Good Z, Chang S, Samusik N, Saumyaa S, Li Y, Zhou X, Liang R et al (2022) Identification of cell types in multiplexed in situ images by combining protein expression and spatial information using CELESTA. Nat Methods 19: 759–769

Zhou BO, Yue R, Murphy MM, Peyer JG, Morrison SJ (2014) Leptin-receptor-expressing mesenchymal stromal cells represent the main source of bone formed by adult bone marrow. Cell Stem Cell 15: 154–168

Zhou Y, He W, Hou W, Zhu Y (2024) Pianno: a probabilistic framework automating semantic annotation for spatial transcriptomics. Nat Commun 15: 2848

